# Kinetochore and ionomic adaptation to whole genome duplication

**DOI:** 10.1101/2023.09.27.559727

**Authors:** Sian M. Bray, Tuomas Hämälä, Min Zhou, Silvia Busoms, Sina Fischer, Stuart D. Desjardins, Terezie Mandáková, Chris Moore, Thomas C. Mathers, Laura Cowan, Patrick Monnahan, Jordan Koch, Eva M. Wolf, Martin A. Lysak, Filip Kolar, James D. Higgins, Marcus A. Koch, Levi Yant

## Abstract

Transforming genomic and cellular landscapes in a single generation, whole genome duplication (WGD) brings fundamental challenges, but is also associated with diversification. How is WGD tolerated, and what processes commonly evolve to stabilize the resulting polyploid? Here we study this in *Cochlearia* spp., which have experienced multiple WGDs in the last 300,000 years. We first generate a chromosome-scale genome and sequence 113 individuals from 33 diploid, tetraploid, hexaploid, and outgroup populations. We detect the clearest post-WGD selection signatures in functionally interacting kinetochore components and ion transporters. We structurally model these derived selected alleles, identifying striking WGD-relevant functional variation, and then compare these results to independent recent post-WGD selection in *Arabidopsis arenosa* and *Cardamine amara*. Most prominent in these results is genetic evidence of at least four functionally interacting kinetochore complex subunits in adaptation to WGD at the centromere among our very top selective sweep outliers. In addition, some of the same biological processes evolve in all three WGDs, but specific genes recruited are flexible. This points to a polygenic basis for modifying systems that control the kinetochore, meiotic crossover number, DNA repair, ion homeostasis, and cell cycle. Given that DNA management (especially repair) is the most salient category with the strongest selection signal, we speculate that the generation rate of structural genomic variants may be altered by WGD in young polyploids, contributing to their occasionally spectacular adaptability observed across kingdoms.

**Significance Statement:** Whole-genome duplication (WGD) occurs in all kingdoms and is linked to adaptation, speciation, domestication, and even cancer outcome. But WGD is a shock to the system, and commonly disrupts cell division due to increased DNA management burden and transformed cell physiology. Nevertheless, the hopeful monster that survives WGD is special, occasionally experiencing runaway success. Why do some thrive but others die? Here we introduce a powerful new model, Cochlearia, which has benefitted from multiple WGDs, and we provide the first genetic evidence of rapid adaptation of functionally interacting components of the cell division machinery, the kinetochore. We also compare which processes and genes evolve to stabilize the new polyploid in three independent cases and highlight common mechanisms.

## Introduction

Whole genome duplication (WGD, leading to polyploidy) is a dramatic mutation that disrupts fundamental cellular processes. Yet, for those that can adapt to a transformed polyploid state, WGD holds great promise^1–3^. Despite its importance to evolution, agriculture, and human health, we do not yet know why some polyploids thrive, while others do not^1,4^.

Immediately following WGD in autopolyploids (within-species WGD, not hybrid allopolyploids), novel challenges arise. The most obvious concerns meiosis: the instant doubling of homologs complicates chromosome pairing^5^. If a chromosome engages in crossing over with more than one homolog, entanglements and breakage ensue at anaphase. WGD also disrupts cellular equilibria, including cell division, ion homeostasis and cell size^6^. These challenges are insurmountable for many nascent polyploids, although established autopolyploids can persist, indicating that they can be overcome.

To date, work in two diploid-autotetraploid model systems has explicitly sought a basis of adaptation to WGD in recent (< 300,000 year-old) autopolyploids. In *Arabidopsis arenosa,* a handful of physically and functionally interacting meiosis proteins undergo adaptive evolution less than 200,000 years post-WGD^7,8^. Derived alleles of these genes decrease chromosome crossover rates, stabilizing meiosis^9–12^. Next, a pool-seq-based scan in *Cardamine*, a genus 17 million years diverged from *Arabidopsis,* showed only very modest convergence with *A. arenosa* on the level of functional pathways for processes under selection shortly post-WGD^13^. Meiosis showed little convergence and the signal of evolution of the meiosis genes obviously controlling crossovers seen in *A. arenosa* was absent. These works gave insight but had important limitations: there was very low sample number in *A. arenosa* (24 individuals^7^), and in *C. amara,* pool-seq of only four populations using a highly fragmented reference^13^. More significantly, as noted in the *C. amara* study, widespread vegetative reproduction in *C. amara* offers some escape from selection for meiotic stability post-WGD, consistent with the results^13^. Thus, minimal convergence between these systems leaves unresolved what salient processes stabilize recent polyploids. This is important because the genomic changes that occur post-WGD may also help explain why some polyploids are so successful and most are not.

Here we address this in a novel system that overcomes these limitations in a more distantly related, independent, and successful set of WGD events within a single species flock (at least 5 WGDs in *Cochlearia* the last 300,000 years)^14^. The *Cochlearia* species complex exhibits diploid, autotetraploid, allohexaploid, octoploid and heptaploid cytotypes (Fig.1A)^14–17^, with the widespread autotetraploid cytotype^14^, similar in age to *A. arenosa*^18,19^. *Cochlearia* is found across Europe, from Spain to the Arctic, in a wide range of habitats including freshwater springs, coastal cliffs, sand dunes, salt marshes, metal contaminated sites and roadside grit (Fig. 1A)^14,20–30^. A broad habitat differentiation is evident by ploidy, with diploids typically found in upland freshwater springs, autotetraploids overwhelmingly on coasts, often directly adjacent to seawater or continuously submerged, and hexaploids in extreme salt marsh conditions. In fact, the hexaploid *Cochlearia danica* is one of the most rapidly spreading invasive species in Europe, invading salted roadways since the 1970’s^23,31^.

**Figure 1.**
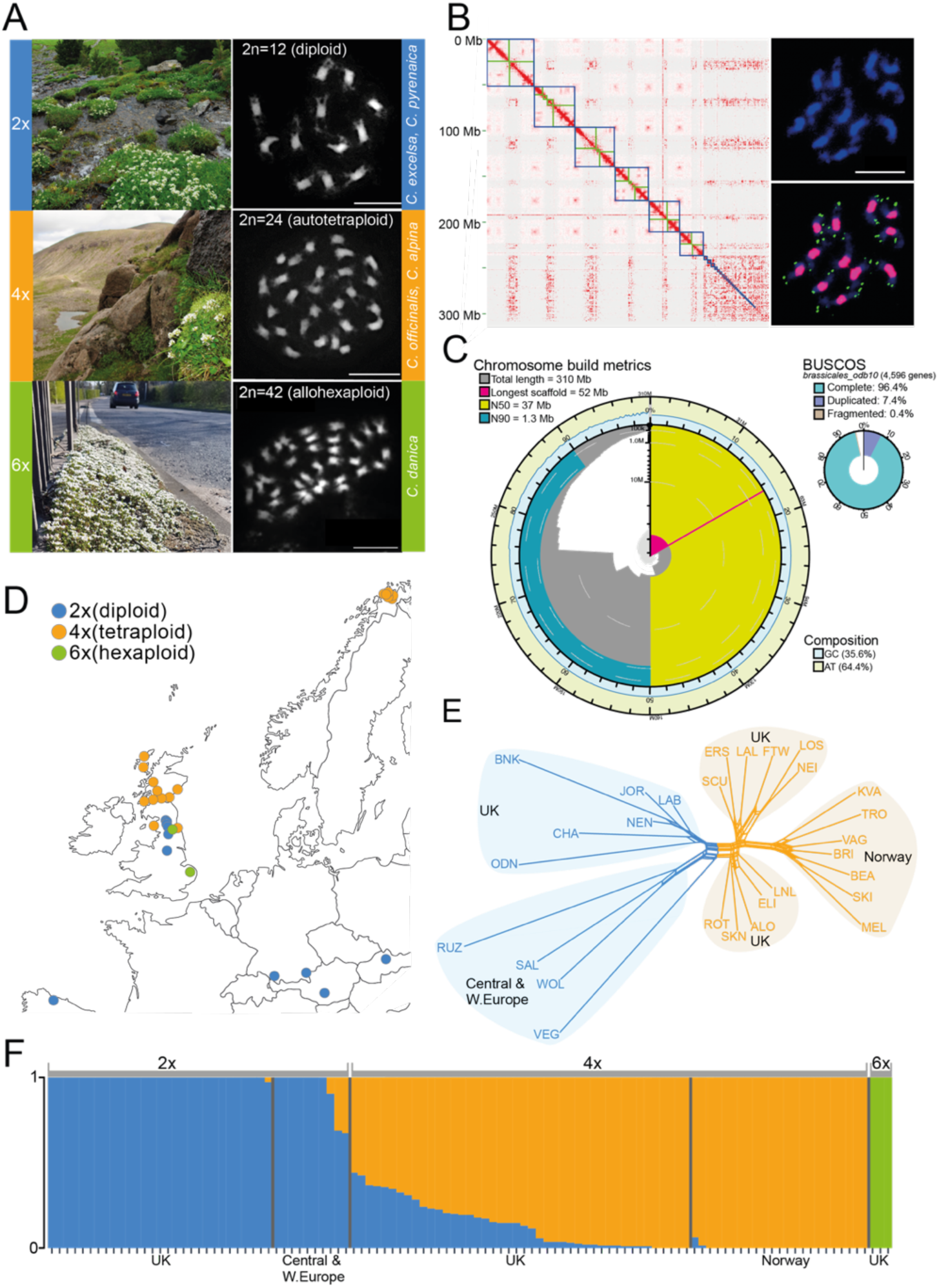
Ploidy variation, genome assembly, sampling, and genetic structure. **(A)** Three *Cochlearia* species in this study (top: *C. pyrenaica* [2n=2x=12], middle: *C. officinalis* [2n=4x=24], bottom: *C. danica* [2n=6x=42]. Scale bars, 10 μm; **(B)** Hi-C contact map and cytology used to orient chromosome arms. Chromosomes are bounded in blue and centromeres indicated in green. Cytology shows DAPI-stained chromosomes with heterochromatic pericentromeres (top) and 102 bp satellite (pink) and Arabidopsis-type telomeric satellite (green) probes hybridizing to all (peri)centromeres and telomeres, respectively (bottom). Scale bar, 10 μm; **(C)** Chromosome-scale assembly of the diploid *C. excelsa* genome; **(D)** The 30 *Cochlearia* populations included in the short read sequencing (locations in Dataset S2); **(E)** Nei’s genetic distances between the populations (hexaploids excluded) visualized in SplitsTree^39^; **(F)** fastSTRUCTURE^40^ analysis (*k*=3, min alleles=8) of all *Cochlearia* individuals in the study, with regions and ploidies indicated. Note that the same color legend applies to panels A, D, E and F: Blue=diploids; orange=tetraploids; green=hexaploids.

We first assess *Cochlearia* demography by individually resequencing 113 plants from 33 diploid, autotetraploid, hexaploid, and outgroup populations from across its ploidy-variable range. We then focus our analysis on closely related diploids and autotetraploids in the UK and scan for selective sweeps post-WGD. We dissect functional targets of adaptive evolution post-WGD using protein modelling and identification of orthologous derived sites from functional studies. Our results show convergence at the process level in very recent WGD adaptation events in these three genera separated by greater spans of ∼40 million years. This indicates that similar processes adapt in response to WGD, but that specific genes recruited are far less constrained. Surprisingly, also we also find strong signal of post-WGD evolution in several kinetochore components, pointing to a novel mechanism of adaptation to polyploid meiosis and mitosis.

## Results and Discussion

### Chromosome-level assembly

To serve as a reference for our demographic analysis and selection scans, we built a chromosome-level genome assembly of one diploid *Cochlearia excelsa* individual (Styria, Gurktaler Alpen, Austria). We chose *C. excelsa* because it is an early diverging diploid (2n = 12) and conveniently also a rare primarily selfing *Cochlearia* species. This resulted in a highly contiguous primary assembly (contig N50=15 Mb; Fig.1B), generated from Oxford Nanopore PromethION data (read N50=27 Kb). The primary assembly was performed with Flye 2.9^32^ and NECAT^33^ with one round of polishing in Medaka^34^ and Pilon^35^ The assembled contigs were then scaffolded to chromosome scale using Hi-C (Fig.1B) and a final cleanup performed with Blobtools^36^ (Fig. S1). Hi-C-guided chromosome arm orientations were confirmed with concordant FISH and in silico mapping of telomeric and centromeric repeats (Fig.1B; Fig. S2). This assembly consists of six primary scaffolds corresponding to the six *C. excelsa* chromosomes with a scaffold N50 of 37 Mb and an overall genome size of 310 Mb (Fig.1B, C), matching our estimated haploid genome size of 302 Mb (Fig. S3). Gene space representation was very good, with 96.4% complete Brassicales BUSCOs^37^ found in the assembly Fig.1C). We performed an annotation incorporating RNA-seq data from the reference line (flower bud, leaf, stem, and silique), protein homology information, and *ab initio* modelling with BRAKER2^38^. This yielded 54,424 gene models across the six chromosomes.

### Geographic distribution, ploidy variation, cohort construction and population sequencing

To determine optimal population contrasts for WGD-specific signatures of selection, we sampled populations across the reported range of the *Cochlearia* species throughout Europe^14,17,22–25,27,28,30^ and conducted flow cytometry- and cytologically-based surveys of genome size and ploidy variation (Fig.1A; Dataset S1 Ploidy Survey; Fig. S4). Measurements were normalised against the diploid population with the most stable individual within-population genome size estimates, WOL (Dataset S1). We focus our demographic analysis on the three most abundant ploidies (Fig.1A): diploids = *Cochlearia pyrenaica* (inland UK and mainland Europe), autotetraploids = *Cochlearia officinalis* (coastal UK and Norway), hexaploids = *Cochlearia danica* (inland UK). Based on these ploidy surveys we chose 113 individuals from 33 populations for sequencing by Illumina PE (average per-individual depth = 17x; minimum = 4x; Fig.1D; Dataset S2 Sample Metrics). The final dataset consisted of 18,307,309 SNPs, on average one variant every 17 bp (quality and depth filtered; Methods).

### Genetic structure

To assess structure in our dataset, we first performed fastSTRUCTURE^40^ (Fig. 1F) on our 109 *Cochlearia* individuals, excluding outgroup sister genus Ionopsidium (23,733 LD-pruned, biallelic 4-fold-degenerate SNPs; max 20% missing data; min minor allele frequency = 0.02). *K*=3 maximized marginal likelihood and grouped samples by ploidy. Focusing on diploids vs tetraploids, PCA confirmed that ploidy dominates over geography (PC1 [ploidy] = 27% of variance explained; PC2 [UK vs Norwegian tetraploids] = 11% of variance explained; Fig. S5). This is reflected also in clean geographical groupings by SplitsTree^39^ analyses, which visualize simple genetic distances (Fig. 1E). This is consistent with a rapid inter- or peri-glacial radiation and postglacial migration such as found in other *Cochlearia* species^14^.

### Meiotic and ion homeostasis-related phenotypic shifts upon WGD

In a young autopolyploid, initial maintenance of diploid-like crossover frequencies can lead to chromosome entanglements and breakage at meiosis. We therefore confirmed the establishment of meiotic stability of these autotetraploids (Fig. 2). Similar to *A. arenosa* autotetraploids^7^, we found a significant per-chromosome reduction in class I mature crossovers, evidenced by HEI10 foci (diploids=1.14, tetraploids=1.04; p <0.00001; Mann-Whitney). This translated to meiotic stability in the *Cochlearia* tetraploids, although we observed greater variation in meiotic stability in tetraploids, which varied dramatically both within and between families for multivalent production (Fig. 2C, D). By assessing the number of multivalents, most cells of both ploidies could be characterized as at least ‘medium stability’ (see methods). More diploid lines were ‘highly stable’ and more tetraploid lines had high variability, suggesting segregation for stabilizing factors in the tetraploids (Fig. 2D).

**Figure 2.**
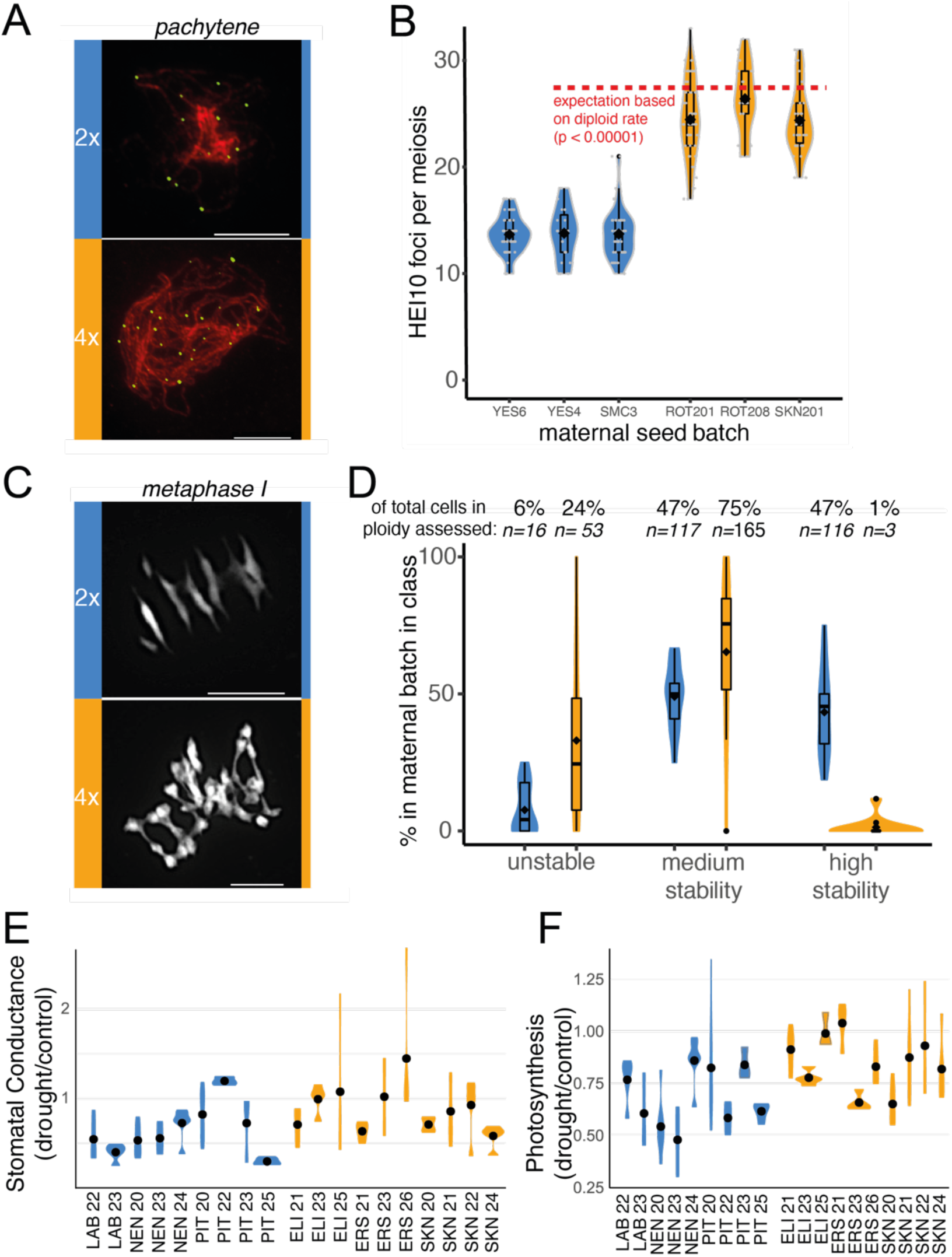
Reduction in crossover number, improved drought response, and high phenotypic variability upon WGD. **(A and B)** Quantification of mature meiotic crossovers by HEI10 staining (green) on pachytene chromosomes stained by ZYP1 (red), showing a significant (p <0.00001; Mann-Whitney) per-chromosome downregulation of crossovers in *Cochlearia* autotetraploids; **(C)** Metaphase I chromosome spreads and **(D)** quantification of meiotic stability, showing decreased, but highly variable stability in tetraploids (orange) relative to diploids (blue). Increased stomatal conductance **(E)** and photosynthetic **(F)** rates under dehydration stress in tetraploids. Plots show median and variation of drought stressed plants in comparison to well-watered plants. (Sign differences between diploids and tetraploids as seen in one-way ANOVA, posthoc Tukey; Table S4) All panels: Blue=diploids; orange=tetraploids. Scale bars, 10 μm.

*Cochlearia* in the UK exhibits a broadly disjunct geographic distribution by ploidy, with diploids (n=6) deeply inland and autotetraploids inhabiting coastal regions of the highest salinity, including full seawater submergence (Fig.1A; Fig. S4). A direct mechanistic link between WGD and salinity tolerance was established in *Arabidopsis thaliana*, where first-generation neoautopolyploids (otherwise isogenic with diploid siblings) show elevated salinity tolerance and intracellular potassium^41^. We therefore tested for ploidy-related differences in salinity tolerance and dehydration stress tolerance in wild *Cochlearia*. Interestingly, in terms of overall plant survival, we found extreme salinity tolerance in all ecotypes tested, with even diploids tolerating up to 600mM NaCl (salinity level of seawater), along with all higher ploidies (Table S1). Tetraploids showed signal of increased drought tolerance, with both elevated stomatal conductance and net photosynthetic rates under drought, relative to diploids (Fig. 2E and F; Tables S2-S4). This suggests an adaptive benefit specific to higher ploidies in response to drought. A benefit under salinity stress may be tempered by preadaptation to a stringent ionic challenge even in the diploids of this species flock, consistent with their halophyte and cold-loving nature^42,43^.

### Selective sweeps associated with WGD

To identify candidate genes and processes mediating adaptation to WGD, we next focused on the 18 geographically proximal populations from the UK (44 autotetraploid individuals and 29 diploid individuals) with good sequencing coverage. Concentrating the selection scan on UK diploids and autotetraploids minimizes the effect of genetic structure that would be introduced if we were to use the mainland European samples as well. To guide our selection scan window size choice, we calculated pairwise linkage decay, which was rapid in both diploids and tetraploids, with near complete lack of genotypic correlations within 2 kb to very low background levels (Fig. 3A). We thus calculated in 1kb windows a battery of differentiation metrics (Dxy^44^, Rho^45^, Hudson’s F_ST_^46^, Nei’s F_ST_^47^, Weir-Cochran’s F_ST_^48^, and groupwise allele frequency difference [AFD]) genome-wide (minimum = 15; mean = 101 SNPs per window). After filtering, these scans overlapped 40,245 of 54,424 total predicted genes, or 74% of gene coding loci with sufficient coverage for assessment.

**Figure 3.**
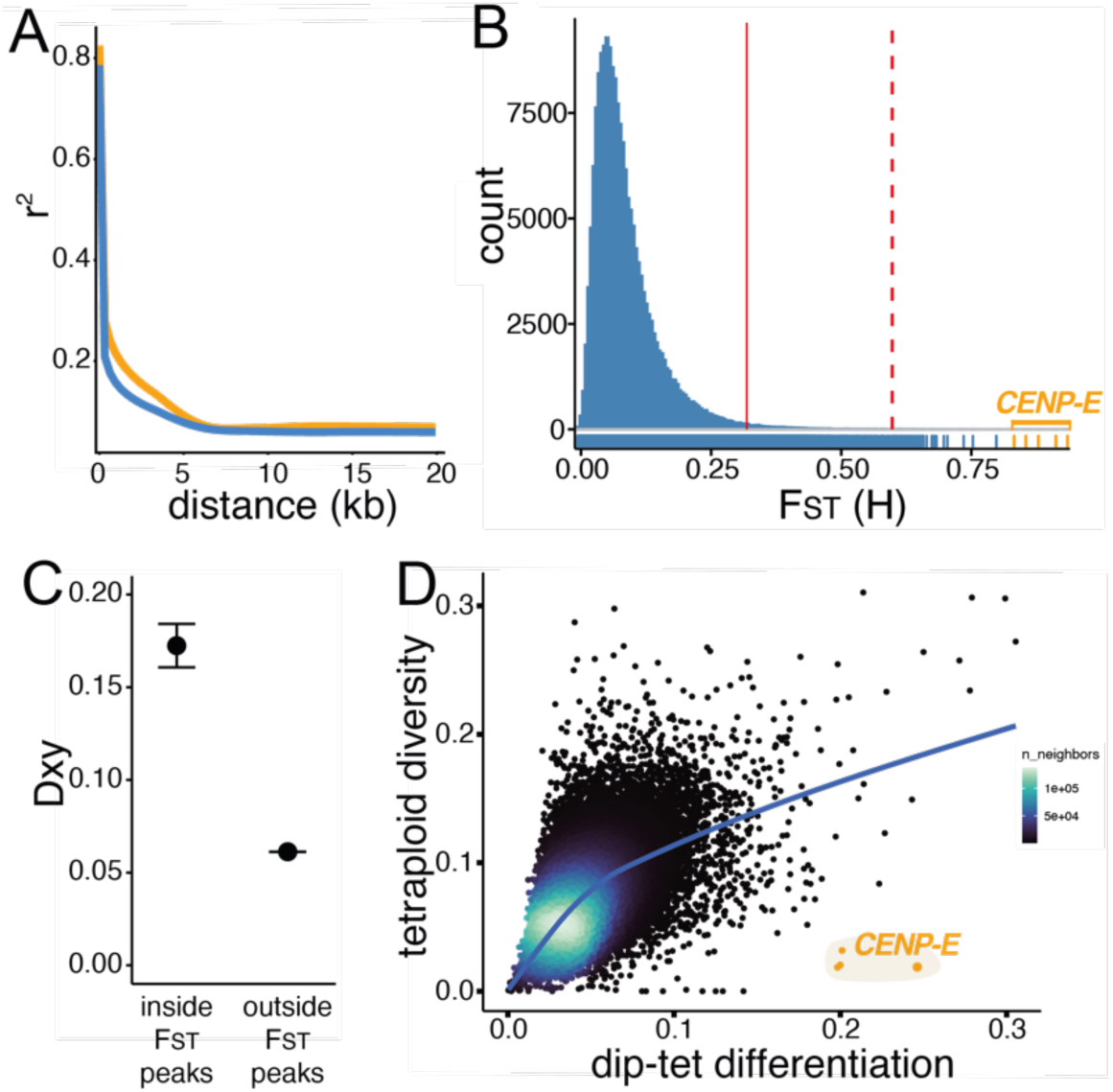
Rapid linkage decay and empirical outlier analysis. **(A)** Immediate decay of genotypic correlations (r^2^) observed in both diploid (blue) and tetraploid (orange) *Cochlearia*; **(B)** Distribution of genome wide F_ST_ values for 182,327 1kb windows. The dashed red line gives the extreme stringency F_ST_ cutoff of the top 25 genome-wide outlier genes; the solid red line gives the 1% cutoff; windows inside the *CENP-E* gene coding region are highlighted in orange; **(C)** Dxy values are significantly elevated inside F_ST_ peaks (Mann Whitney U test: *P* < 2 ξ 10^-16^); **(D)** *CENP-E*, the #1 genome-wide F_ST_ outlier, also exhibits greatly excess differentiation for its level of diversity in the tetraploids, a classical signal of selective sweep.

To empirically determine which metric most reliably identified genomic regions that exhibit localized peaks in AFD indicative of specific sweeps upon WGD, we performed an inspection of all AFD plots in all outlier tails (see Methods). From this, we identified Hudson’s F_ST_ as most reliably identifying regions of localized AFD peaks (and not e.g., low diversity^44^). This formulation of F_ST_ brings the added benefit of robustness for unequal population sizes and presence of rare variants^49^ and direct comparisons to the *C. amara* study, which used this same metric^13^. We therefore extracted windows in the top 1% of this distribution as candidate outliers, consisting of 1,823 1kb windows, overlapping 753 gene-coding loci for which we could obtain functional descriptions primarily from orthology (or, lacking this, close homology) to *Arabidopsis thaliana* (Dataset S3 Selective Sweep Candidates).

We focus on the most extreme 25 of these (to the right of the dashed line in Fig 3B; Table 1), which we confirm exhibit elevated Dxy values (outlier F_ST_ peak Dxy=0.17; mean outside peak Dxy=0.06; Mann Whitney U test: *P* < 2 IJ 10^-16^; Fig. 3C).

**Table 1.** Top selective sweep candidates following WGD in *Cochlearia*. Top 25 of 40,245 genes assessed, with 5 additional genes (bottom) with 10 or more MAV SNPs and in top 1% F_ST_ tail. * = contains 37/39 whole-genome fixed differences between diploids and tetraploids and is also the only gene with three of the top 25 Dxy windows. Genome-wide F_ST_ rank is in Column 1.

| F<sub>ST</sub> rank | Cochlearia ID | A.thaliana ID | Name | Description |
| --- | --- | --- | --- | --- |
| 1* | g7445 | AT3G10180 | CENP-E | **CENTROMERE PROTEIN E.** Kinetochore protein that moves mono-oriented chromosomes to the spindle equator<sup>51</sup>. Cooperates with chromokinesins and dynein to mediate chromosome congression. Activity is regulated by post-translational modifications, protein interactions and autoinhibition. Tetraploid cancers are far more susceptible to CENP-E inhibitors than diploids<sup>52-54</sup>. |
| 2 | g31016 | AT1G06670 | NIH | NUCLEAR DEIH-BOX HELICASE. Binds DNA without clear specificity<sup>55</sup>. |
| 3 | g7446 | AT3G10070 | TAF12 | TATA-ASSOCIATED II 58. Controls stress responsive root growth<sup>56</sup> |
| 4 | g40185 | AT2G26690 | NPF6.2 | NITRATE TRANSPORTER 6.2. Mediates drought stress response<sup>57</sup> |
| 5 | g25338 | AT4G00060 | MEE44 | MATERNAL EFFECT EMBRO ARREST 44: Part of the RNA TRAMP complex |
| 6 | g49945 | AT1G15660 | CENP-C | **CENTROMERE PROTEIN C.** Kinetochore protein that is critical for centromere identity in both mitosis and meiosis. Loss of CENP-C results in aneuploidy and cell death<sup>60</sup>. Necessary in mitotic cells for kinetochore assembly centromere establishment. Mutants also fail to retain centromeric SC in late pachytene<sup>58,59</sup>. |
| 7 | g25334 | AT1G05940 | CAT9 | CATIONIC AMINO ACID TRANSPORTER 9 |
| 8 | g25335 | AT4G00026 | SD3 | **SEGREGATION DISTORTION 3.** Mutants have lower ploidy<sup>60</sup>. |
| 9 | g6996 | AT3G16730 | CAP-H2 / HEB2 | **CONDENSIN-II COMPLEX SUBUNIT H2.** Functions in DSB repair and arranges interphase chromatin<sup>61,62</sup>. Plays a role in alleviating DNA damage and genome integrity in *A. thaliana*<sup>62</sup>. Condensin II-depleted human cells have a defect in homologous recombination-mediated repair<sup>63</sup>. |
| 10 | g24649 | AT5G40595 |  | Unknown protein |
| 11 | g24648 | AT4G02000 |  | Ta11-like non-LTR retrotransposon |
| 12 | g33330 | AT1G54310 |  | S-adenosyl-L-methionine-dependent methyltransferases superfamily protein |
| 13 | g40302 | AT4G10310 | HKT1 | **HIGH-AFFINITY K<sup>+</sup> TRANSPORTER 1.** Sodium transporter. Mediates salinity tolerance in wild *A. thaliana* populations<sup>64</sup>. Under selection post-WGD in *A. arenosa*<sup>7</sup>. |
| 14 | g33311 | AT5G61390 | NEN2 | NAC45/86-DEPENDENT EXONUCLEASE-DOMAIN PROTEIN 2 |
| 15 | g10739 | AT2G01980 | SOS1 | **SALT OVERLY SENSITIVE 1.** Plasma membrane-localized Na<sup>+</sup>/H<sup>+</sup> antiporter that extrudes Na<sup>+</sup> from cells. Is essential for plant salt and stress tolerance. |
| 16 | g46402 | AT1G77990 | SULTR2 | SULPHATE TRANSPORTER 2;2. A low-affinity sulfate transporter |
| 17 | g4631 | AT5G19270 |  | Reverse transcriptase-like protein |
| 18 | g4632 | AT2G07200 |  | Cysteine proteinases superfamily protein |
| 19 | g25121 | AT5G35970 |  | P-loop containing nucleoside triphosphate hydrolases superfamily protein |
| 20 | g45503 | AT5G08620 | STRS2 | STRESS RESPONSE SUPPRESSOR 2. A DEA(D/H)-box helicase involved in drought, salt and cold stress responses. |
| 21 | g25112 | AT5G35980 | YAK1 | YAK1-related. Controls cell cycle and regulates drought tolerance<sup>65,66</sup>. |
| 22 | g40349 | AT4G11110 | SPA2 | SPA1-related 2. Convergent with WGD adaptation in *A. arenosa*<sup>7</sup>. |
| 23 | g54387 | AT3G01420 | DOX1 | An alpha-dioxygenase involved in protection against oxidative stress. |
| 24 | g25300 | AT5G34940 | GUS3 | GLUCURONIDASE 3 |
| 25 | g45090 | AT1G65320 | CBSX6 | Cystathionine beta-synthase family protein |
| 15 mav | g39361 | AT5G55820 | INCEP / WYRD | **INNER CENTROMERE PROTEIN.** The largest subunit of the Chromosome Passenger Complex (CPC), and directly binds to all other subunits in animals and yeast<sup>67</sup>. The CPC ensures that all kinetochores are attached to microtubules emanating from opposing poles. INCENP is necessary for normal mitotic divisions<sup>68</sup>. At mitosis and meiosis localises to kinetochores, and later, the phragmoplast<sup>67</sup>. |
| 12 mav | g25323 | AT4G29090 |  | Ribonuclease H-like superfamily protein |
| 11 mav | g23778 | AT4G25290 |  | DNA photolyase |
| 11 mav | g54385 | AT1G56145 | CORK1 | A LRR receptor kinase required for celooligomer-induced responses. |
| 10 mav | g33400 | AT1G50240 | FU | **FUSED.** An ARM repeat domain-containing protein kinase involved in male meiosis. Tightly localized to nascent phragmoplast and with the expanding phragmoplast ring. |

Complementing this approach, we also focused our 1% outlier list on gene coding regions with a fineMAV approach^50^. Using Grantham scores to estimate functional impact of each non-synonymous amino acid change encoded by a given SNP, this approach scales the severity of predicted amino acid change by the AFD between groups. Of the 107,055 non-synonymous-encoding SNPs assigned a MAV score, the top 1% outliers from the empirical distribution were intersected with our F_ST_ outliers, yielding a protein-evolution-oriented list of 159 gene coding loci, harboring 290 MAV SNPs (bold in Dataset S3 Selective Sweep Candidates; 1% F_ST_ outliers with 10 or more 1% extreme MAV outliers are given in Table 1). By these approaches, we could resolve clear gene-specific peaks of F_ST_ (Fig. 4A) and candidate selective sweep alleles in our top 25 genome-wide outliers (Fig. 4 B-G).

**Figure 4.**
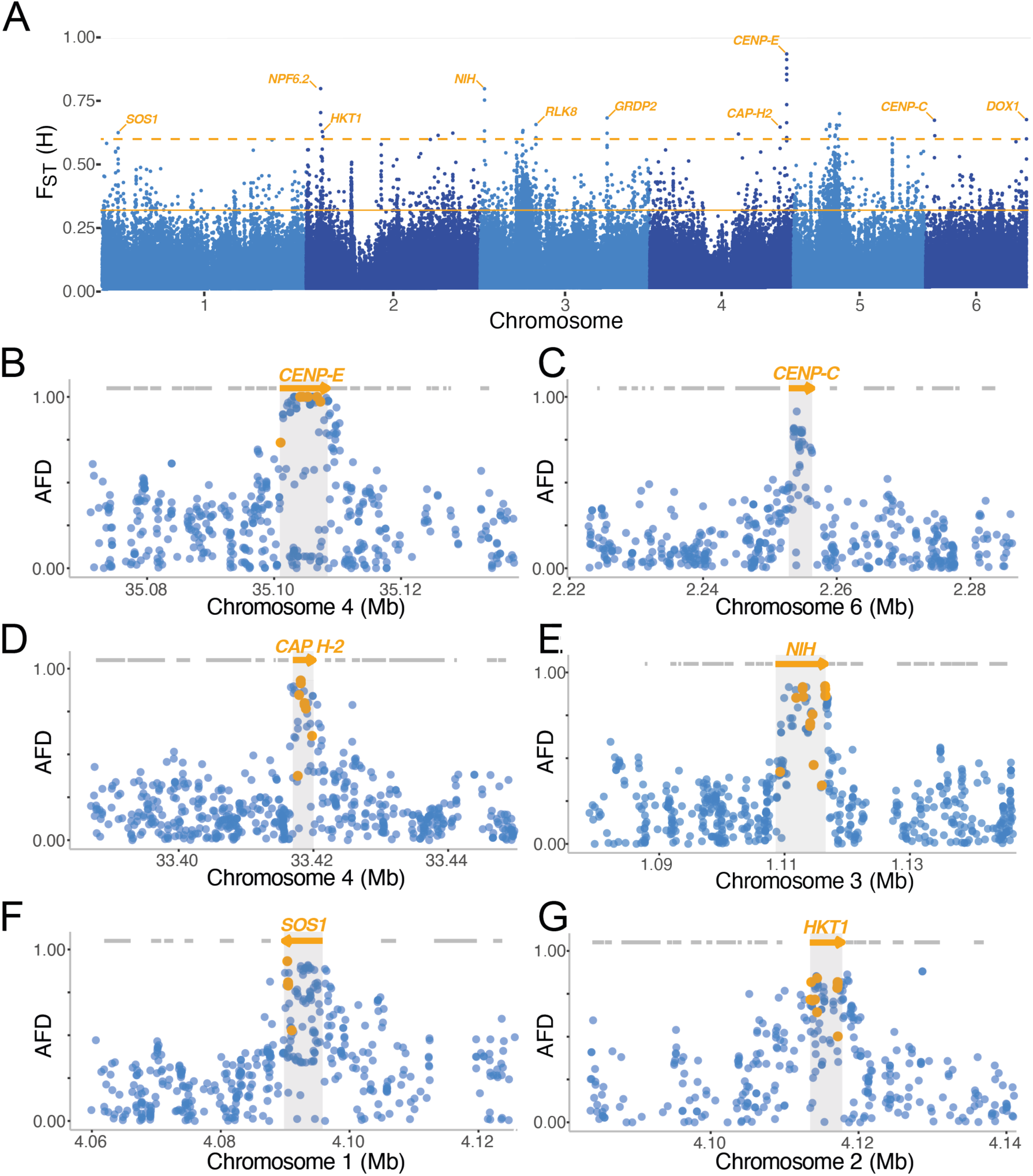
Selective sweep signatures of DNA management and ion homeostasis alleles. **(A)** Ploidy-specific differentiation across the *Cochlearia* genome. The dashed orange line gives the extreme stringency F_ST_ cutoff of the top 25 outlier genes (of 40,245 assessed); the solid line gives a 1% F_ST_ cutoff; **(B-G)** Examples of selective sweep signal among 6 top outlier genes; **(B-E)** Represent kinetochore or DNA management; **(F-G)** ion homeostasis functional categories. The X axis gives genome position in megabases (Mb). The Y axis gives AFD values at single SNPs (dots) between diploid and tetraploid *Cochlearia*. Orange arrows indicate genes overlapping the top F_ST_ gene outliers, and grey lines indicate neighboring genes. Orange dots indicate MAV outliers.

### Functional processes under selection post-WGD in *Cochlearia*

While we focus our discussion below to the top 25 genome-wide outliers, a broader, 1% F_ST_ list of 753 selective sweep candidates (Dataset S3) yield particularly informative gene ontology (GO) enrichments, with 181 significantly enriched categories (using a conservative ‘elim’ Exact Test; Dataset S4 Gene Ontology Enrichment Cochlearia). Many of the enriched categories can be grouped into three classes congruent with WGD-associated changes^1,2,4–7,69^: 1) DNA management: 29 categories relate to DNA integration, cell division, meiotic chromosome segregation, mitosis, DNA repair, and recombination; 2) Ion homeostasis: 26 categories relate to ion transport (principally extrusion), cation homeostasis, salt stress, and stomata; and 3) Cell stoichiometry: 7 categories relate to global gene expression, cell wall, and biosynthetics, pointing to both global gene expression and ‘nucleotypic’^6^ changes upon WGD. These categories often overlap, for example ‘cell stoichiometry’ and ‘DNA management’ in terms of global RNA transcriptional changes post-WGD, which should alter due to doubled DNA template.

### Kinetochore evolution upon WGD

Genome-wide, the most dramatic selection signature is directly over the coding region of the *CENTROMERE PROTEIN E* ortholog (*CENP-E*; Fig. 4A, B)^70^. This gene overlaps the 5 top F_ST_ outlier windows (mean F_ST_ = 0.88) and includes 37 of the 39 genome-wide fixed differences between ploidies. An essential kinetochore protein, *CENP-E* moves mono-oriented chromosomes to the spindle equator, mediating congression^51,71,72^. Strikingly, tetraploid cancers are far more susceptible to CENP-E inhibitors than diploids^52–54,73,74^.

The *Cochlearia* CENP-E coding region contains 26 SNPs (synonymous or non-synonymous) that are highly differentiated between diploids and tetraploids (>50% AFD). Six of these are unique to *Cochlearia* tetraploids, at highly conserved sites across angiosperms (Fig. 5A). None are in characterized conserved functional regions of the kinase domain^75^, meaning motor activity is likely intact. Most of the tetraploid-specific changes are in the coiled coil regions, which in animals are important for regulation of cell division via phosphorylation and protein-protein interactions^51^. For example, point mutations in the coiled coils are associated with human disease (e.g., microcephaly^76,77^). In humans and *Xenopus*, these regions are known to be extensively phosphorylated during the cell cycle and may be involved in the autoinhibition of CENP-E^78–81^. Indeed, we see four tetraploid-specific changes that may affect regulation via the loss of phosphorylation (S717A, S821A, S1059L and S1169). Three additional changes toward the C-terminus are in a cargo (chromosomes) binding region. Four of the tetraploid-specific changes show remarkable conservation across plants, being otherwise absolutely conserved across CENP-E-like kinesins (Fig. 5A; A607G, Q613D, R899Q and Q1024E). Taken together, these data suggest changed regulation of CENP-E at mitosis and/or meiosis. This is consistent with functional evidence from *A. thaliana* showing that *CENP-E* mutations extend the cell cycle^81^.

**Figure 5.**
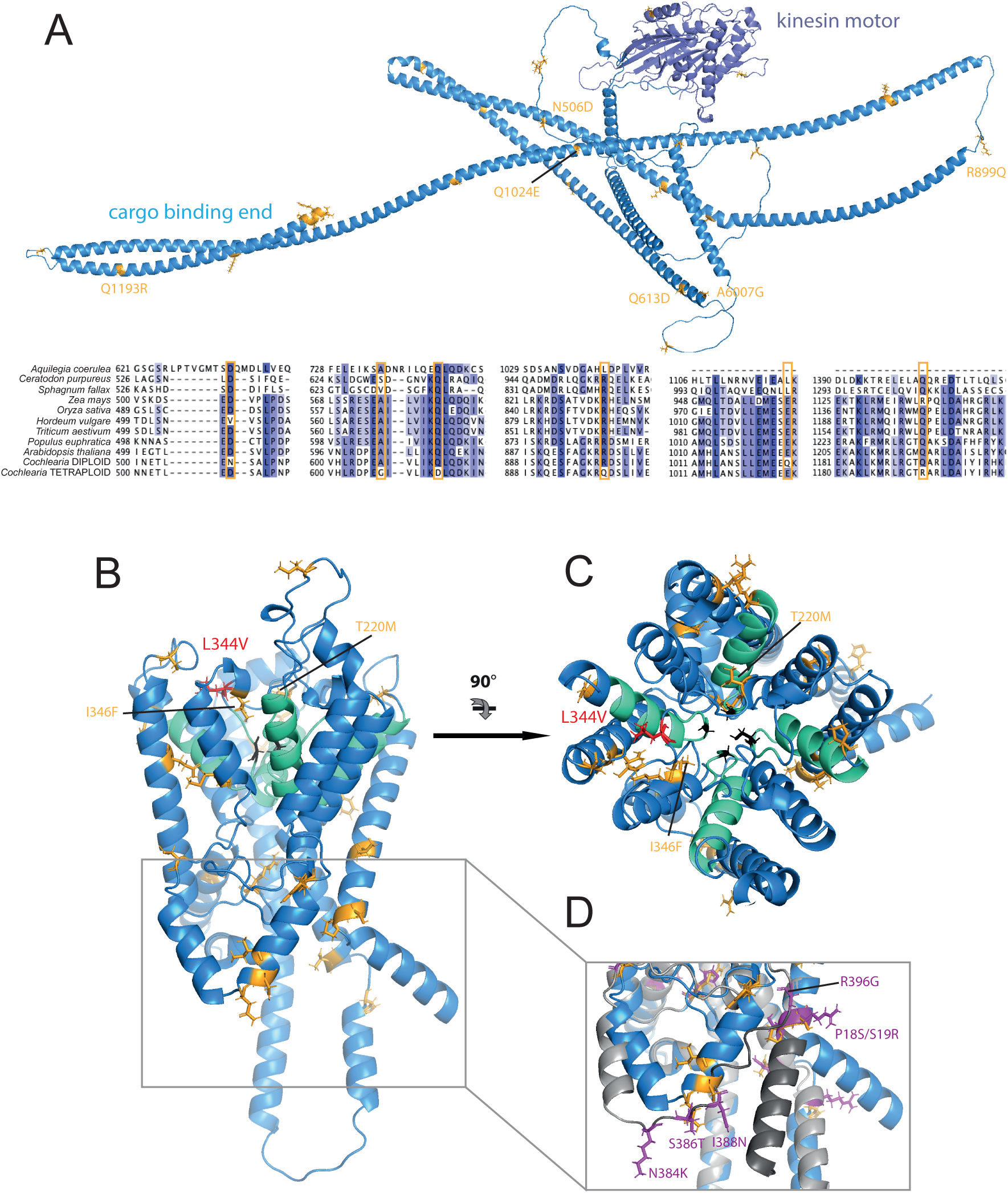
Tetraploid-derived protein structure changes in CENP-E and HKT1. **(A)** Kinetochore subunit CENP-E structural prediction and pan-plant diversity alignment. Structure: diploid structure (blue), consensus (>50% allele frequency) derived polymorphisms in tetraploids (orange), kinesin motor domain (slate). Alignment: colour by percentage identity; consensus mutations at highly conserved sites (orange boxes); **(B-D)** Ion channel HKT1: diploid structure (blue), mutation sites (orange; diploids, magenta: tetraploids), pore domains (teal); **(C)** HKT1 extracellular surface. Salinity tolerance-mediating change L344V in red and selectivity filter residues S/G1, G2, G3 and G4 (black); **(D)** superimposition of diploid (blue) and tetraploid (grey) intracellular domains. Structures that are predicted to have rotated are highlighted on the tetraploid structure (dark grey). Tetraploid consensus mutations (magenta).

Our top outlier list (Table 1) contains two additional orthologs of *A. thaliana* kinetochore components: *CENP-C*, an essential kinetochore component in both mitosis and meiosis, needed for centromere identity in plants, yeast, *Drosophila*, and humans^58,59,82^ (Fig. 4C) and *INNER CENTROMERE PROTEIN (INCENP)*, which controls mitotic and meiotic chromosome segregation and cytokinesis in plants, yeast and animals^68,83^. At mitosis and meiosis *INCENP* localizes to kinetochores, and later, the phragmoplast as the main subunit of the Chromosome Passenger Complex^67^. Both INCENP and CENP-C contain 1% outlier-MAV SNPs, with *INCENP* harboring a remarkable 15 MAV outlier SNPs, the greatest number of any gene in the genome.

### Evolution of DNA repair and transcription

Several of the top signals of selective sweeps are in DNA repair-related genes, for example in *CONDENSIN-II COMPLEX SUBUNIT H2/HYPERSENSITIVITY TO EXCESS BORON 2* (Fig. 4D), which functions directly in double-strand break (DSB) repair^62,84^ and chromatin management in plants, mouse and *Drosophila*^61,85^. *Condensin II* is required for proper DNA DSB repair by homologous recombination (HR) repair in *A. thaliana* and humans^62,63^, and has been implicated association and dissociation of centromeres^85^. We find in our 1% F_ST_ outliers an outlier also at a homolog of *DAYSLEEPER*, an essential domesticated transposase^86^. *DAYSLEEPER* binds the Kubox1 motif upstream of the DNA repair gene *Ku70* to regulate non-homologous end joining double-strand break repair, the only alternative to HR^87,88^.

In a young polyploid, the total DNA content doubles but the protein content and cell size does not scale accordingly^6^, so we predicted that the control of global gene expression, like meiosis, should undergo adaptive evolution post-WGD. Here we see signal of this, with a suite of DNA or RNA polymerase-associated genes among our selective sweep outliers. In our 1% F_ST_ outliers this includes *NRPB9*, an RNA polymerase subunit that is implicated in transcription initiation, processivity, fidelity, proofreading and DNA repair^89–93^ as well as the ortholog of *MED13*, of the mediator complex, which is essential for the production of nearly all cellular transcripts^94^ (Dataset S3).

### Evolution of ion homeostasis, transport and stress signaling

The ionomic equilibrium of the cell is immediately disrupted at WGD^41^; in particular K^+^ concentrations are increased instantly, consistent with increases in salinity tolerance in synthetic *A. thaliana* autotetraploids^41^. In our young tetraploids, among the top selective sweeps are ion channels that function explicitly to remove K^+^, Na^+^ and other cations from the cell. At F_ST_ rank 13 genome-wide, we see the ortholog of *HIGH-AFFINITY K^+^ TRANSPORTER*^64,95^ *(HKT1;* Fig 4G, 5B), and at rank 15, *SALT OVERLY SENSITIVE 1 (SOS1)*, a membrane Na^+^/H^+^ transporter that removes excessive Na^+^ from the cell and is central to salt tolerance^96,97^ (Fig 4F). Several studies have demonstrated adaptive natural variation in response to salinity in *HKT1*^64,95^, but among natural systems there have been to date no works until now that implicate *SOS1,* although this gene is central to salt tolerance pathways (discussed in ^98^).

Structural homology modeling confirms that the *Cochlearia HKT* under selection is class 1 from its selectivity filter residue configuration (S-G-G-G), indicating it is likely Na^+^ selective (and K^+^ non-selective)^99^. There is a L344V mutation in the tetraploid relative to the diploid; remarkably, this is the identical site and amino acid change that is associated with salt tolerance in rice OsHKT1;5 (Fig 5B)^100,101^: functional confirmation in rice shows the orthologous site substitution to valine in our tetraploid to be associated with salt tolerance (including faster Na^+^ transport), while the diploid leucine is associated with salt sensitivity (including slower Na^+^ transport)^100^. Given that the tetraploids live overwhelmingly in coastal regions where they are exposed to extreme Na levels, while the diploid lives in low Na freshwater streams, this makes biological sense. While the close proximity of L344 alone is likely enough to disrupt pore rigidity via its larger side chain relative to V, in *Cochlearia* HKT1 we also see T220M (a highly conserved residue) and I346F mutations which are even closer to the P1-4 pore domains that hold S-G2-G3-G4 together to create the selectivity control. These mutations all introduce large side chains that likely affect structural dynamics at the P1-4 pore^100,101^. In addition, mutations F326I, R200P L180F, Q303H, M360I are all in (R200P and F326I) or in contact with the four alpha helices that stabilize the SGGG selectivity filter. While this may seem an excessive quantity of mutations, suggesting gene inactivation and relaxation of selection, all the sites except M360I, T220M and L344V are loosly conserved, suggesting flexibility in these regions.

There are also mutations on the cytosolic side of the protein, a few poorly conserved residues appear to induce small structural changes (Fig 5D). This includes a cluster of changed residues (I388N, S386T, N384K, R396G) which are predicted to break up an alpha helicase in the tetraploid, and P18S and S19R, which appear to induce a break in the first alpha helicase of the protein. To our knowledge this domain is not functionally characterized, though given this positioning, this could represent a change in signaling or regulation.

Congruent with its conserved central role in ion homeostasis, *SOS1* is highly conserved and tetraploid-specific changes are not near active transport (nucleotide or ion) binding sites, nor dimerization domains. Instead, a tight cluster of 3 mutations marks the boundary between the β-sheet-rich cytoplasmic domain (β-CTD) and the C-terminal autoinhibition (CTA) domain which contains a further 3 mutations (K1014E, T1075S and R1101Q). The CTA is unstructured; however, it has been experimentally shown that truncation, or just two point mutations, can radically change the behavior of the channel, presumably by releasing autoinhibition increasing Na^+^ transport and therefore increasing salt tolerance^102^. Notably, this includes one T1075S substitution: S and T are two of the three amino acids that can be phosphorylated and it has been shown that this exact residue behaves differently when phosphorylated, which suggests that the choice of S or T may have evolutionary consequences^103^.

Ion homeostasis shifts should be associated with changes in responses to salt, osmotic, and cold stress, as all these stressors have a common osmotic basis. Such a link between immediate ionomic changes in the polyploid cell may be a key functional basis for the observed ecotypic differentiation of young polyploids, especially as observed in arctic and alpine conditions. We accordingly see in our top outliers categories of relevant genes, for example among the top candidates the ortholog of *DEAD-BOX RNA HELICASE 25 (STRS2)*, identified in *A. thaliana* as a repressor of stress signaling for salt, osmotic, and cold stress^104,105^. This gene also controls freezing tolerance^106^, highly relevant to the cold-loving arctic and alpine history of *Cochlearia*^14^.

To assess if perhaps these sweeps were better associated with ecotype differences between ploidies, we performed a salt tolerance experiment on diploid and tetraploid plants. Interestingly, given their divergent ecotype preferences (Fig. S4), with tetraploids found in more saline conditions, we found that the diploid *Cochlearia* are in fact more salt tolerant than the tetraploids (p = 2.178 x 10^-05^; See Supplementary Text 1 and Table S5.). This finding also contrasts with observations of increased salinity tolerance in neotetraploid *Arabidopsis thaliana*^41^. Again, however, this may be a signal of preadaptation to osmotic challenge (common to freezing, salinity, and dehydration) across the halophyte *Cochlearia*^42^.

Relevant also to these phenotypes, genes involved in stomatal function were outliers post-WGD, such as the ortholog of *OPEN STOMATA2*, a target of ABA stress signaling to close the stomata during drought response^107^. This gene is an ATPase in the plasma membrane that drives hyperpolarization and initiates stomatal opening^105^. We also see *TOO MANY MOUTHS*, where a mutation leads to disruption of asymmetric cell division during stomata development^108^. Finally, we see selection signal in *STOMAGEN*, which acts on the epidermis to increase stomatal formation^109^. Sweeps in these loci are consistent with the phenotypic shifts we observe of increased stomatal conductance and net photosynthetic rate under drought conditions in tetraploid *Cochlearia* populations relative to diploids (Fig. 2E; Supplementary Text 2).

### Gene-level convergence

To test for convergence at the ortholog level, we first determined orthogroups^110^ between *Cochlearia, A. arenosa,* and *C. amara* (Methods). Top 1% F_ST_ outliers for *Cochlearia* (n=753; Dataset S3), *A. arenosa* (n=452; Dataset S5), and *C. amara* (n=229; Dataset S6) were considered orthologues if they were part of the same orthogroup. By this criterion not a single ortholog was under selection in all three species (Fig. S6A; Dataset S7 Gene Ortholog Convergence). This approach depends on strict 3-way orthogroup assignment, so we then searched for convergence by assigning all genes in the outlier lists to a nearest *A. thaliana* homolog. By this ‘nearest homolog’ criterion, only one gene was selected in all three WGDs: *DAYSLEEPER*, an essential domesticated transposase^86^ with a role in regulating non-homologous end joining double-strand break repair (Fig. S6B; Dataset S8).

Interestingly, by both homolog assignment methods, several of the best *Cochlearia* WGD adaptation candidates are present also in *A. arenosa*: *ASY3,* functionally validated^9,12^ to have an primary role in stabilizing autotetraploid meiosis in *A. arenosa* is also in our 1% F_ST_ outlier list in *Cochlearia*^9^. This gene, along with *ASY1*, is critical for formation of meiotic chromosome axes, and tetraploid alleles of both genes result in fewer deleterious multichromosome associations and more rod-shaped bivalents in metaphase I^12^. We also see *CYCD5;1*, which is a QTL for endoreduplication^111^. Additionally, the salinity and osmotic genes *HKT1* and *OST2* in both top candidate lists (Fig. S6A; Dataset S7 Gene Ortholog Convergence). All of these genes are involved in processes that have been implicated in adaptation to WGD^1,2,4–7^ and therefore stand as good candidates in salient challenges to nascent polyploids. We note that overlap between *Cochlearia* and both *A. arenosa* and *C. amara* candidates was greater than expected by chance (SuperExactTest p=0.0024 and p=0.0047 respectively), but only marginally for *C. amara* and *A. arenosa* (SuperExactTest p=0.014).

### Process-level convergence

We reasoned that there may be similarities in processes under selection between the three independent WGDs, despite modest gene-level convergence. To estimate this, we first compared our GO results from those published in *A. arenosa* and *C. amara*. The much greater signal of overlap (process-level convergence; Fig. S6C) was between *Cochlearia* and *A. arenosa*: of the 113 GO biological process terms significantly enriched in *Cochlearia*, 17 were among the 73 GO terms enriched in *A. arenosa* (Dataset S4). These were high-level GO terms including representatives of ploidy-relevant categories, e.g. ‘cell division’, ‘transmembrane transport, and ‘regulation of RNA metabolic processes’.

Despite this evident convergence, in *Cochlearia* an array of DNA repair and kinetochore genes were among top candidates, signaling a shift relative to *A. arenosa*, where a more focused prophase I-oriented signal emerged primarily around Synaptonemal Complex (SC)-associated proteins mediating lower crossover rates^7^. However, given that *CENP-C* mutants fail to retain an SC at the centromere and *CENP-C* appears to have functions in synapsis, cohesion, and centromere clustering^112^, there may be a closer parallel between the obviously ‘SC-focused’ adaptive response in *A. arenosa* and that in *Cochlearia*. Indeed, aside from *ASY3*, in the *Cochlearia* 1% F_ST_ outlier list we also see *ATAXIA-TELANGIECTASIA MUTATED (ATM;* also with 6 MAV SNPs; Dataset S3). This gene controls meiotic DNA double-strand break formation and recombination and affects synaptonemal complex organization^113^. We also see a *PDS5* cohesion cofactor ortholog and *REC Q MEDIATED INSTABILITY 1* (*RMI1*), which suppresses somatic crossovers and is essential for resolution of meiotic recombination intermediates^114^. Notable also are *BRCA2-like B* (which is essential at meiosis and interacts with *Rad51, Dss1,* and *Dmc1*) and *SHUGOSHIN C* (which protects meiotic centromere cohesion).

Further evidence of functional association outliers found in *Cochlearia* with those in *A. arenosa* can be observed in protein interaction information from the STRING database, which provides an estimate of proteins’ joint contributions to a shared function^115^. Using comparable 1% F_ST_ outlier lists from the two species, we see many connections between these independent WGDs. Here we see particularly large clusters (Fig. 6A,B) in center of the overall network (Fig. 6C). In particular, the endopolyploidy gene^111^ *CYCD5;1,* and *DNA Pol V,* a shared outlier in both species, interact with a broad array of other outliers in each selection scan (Fig. 6A). This analysis reveals, for example, that *DNA Pol V* shares as a top candidate either *NRPB9A* (in *A. arenosa*) or *NRPB9B* (in *Cochlearia*). These subunits are partly redundant interactors with *Pol II, IV*, and *V* and the double mutant is fatal in *A. thaliana*^91^.

**Figure 6.**
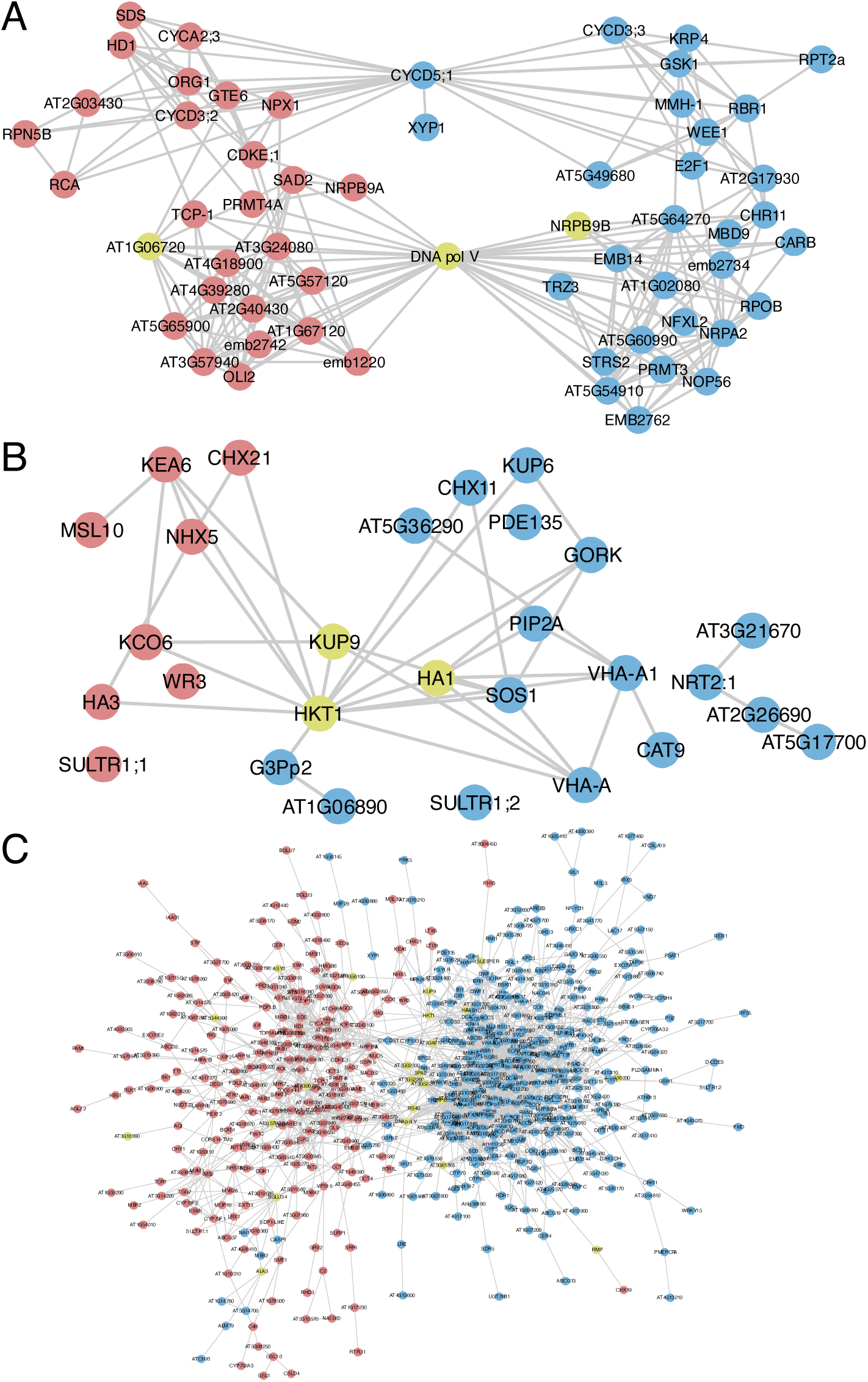
Evidence for functional convergence between *Cochlearia* and *Arabidopsis arenosa* following independent WGDs. STRING^115^ plots show *Cochlearia* candidate genes (blue) and *A. arenosa* candidate genes (red). Convergent genes that are present in both species’ outlier lists as selection candidates are in yellow. **(A)** A large shared cluster surrounding the endopolyploidy gene *CYCD5;1*, which has many connections to large cluster centering on *DNA pol V*, which is an outlier in both datasets; **(B)** ion transport-related genes with a highly interconnected cluster of top outliers in both genome scans, *HKT1, KUP9,* and *HA1;* **(C)** The entire set of candidates in both genome scans for which the STRING database has information.

Taken together, these results indicate that adaptive evolution in response to WGD is focused on particular functions instead of specific genes. These functions involve DNA management and ion homeostasis. However, it is unclear why particular solutions are favored in one species relative to another. A degree of stochasticity depending on available standing variation can be expected, but species histories likely play a role, offering preadaptations that may ‘nudge’ evolution. For example, our analysis of salinity tolerance in *Cochlearia* gave the surprising result that the diploid was at least as tolerant to extreme salt concentrations as the tetraploid, although the diploid is found predominantly inland, except for rare diploid coastal populations from Spain. A postglacial and boreal spread of the diploid towards the UK may have brought salinity and cold tolerance along the way^116^, altering the genomic substrate upon which selection acted in response to WGD-associated ionomic challenge.

## Conclusion

Following whole genome duplication (WGD), the newly polyploid cell requires modifications to chromosome segregation, ion homeostasis, stomatal function, and diverse other processes^1,2,4,6^. Here we investigated the signals of adaptive evolution post-WGD in a successful novel polyploid system, *Cochlearia*. We discovered striking signals of selection in core kinetochore components, *CENP-E, CENP-C, INCEP, CAP-H2,* and others, as well as well-studied ion homeostasis loci, *HKT1* and *SOS1.* We detail specific changes in these proteins upon WGD which are known in model systems to directly modulate WGD-relevant function. We also compare our results to independent WGD adaptation events, finding convergence of these processes, but not genes, indicating a highly flexible array of adaptive mechanisms.

Our results also suggest a hypothesis for the occasionally spectacular adaptability of polyploids. We observed a broad array of DNA management and repair processes under selection in all species, and especially *Cochlearia*. This may signal a temporarily increased post-WGD susceptibility to DNA damage, due to suboptimal function of DNA repair genes during the process of adaptation to the WGD state. This may result in a relative ‘mutator phenotype’ in neopolyploids. Such a mutator phenotype has been plainly observed in polyploid metastatic human cancers, which not only exhibit SNP-level hypermutator phenotypes, but also dramatic structural variation in malignant aneuploid swarms that are associated with progression^3^. We speculate that a parallel to this may exists following other WGDs. Whether this hypothesis is further supported by future discoveries, the centrality of WGD to evolution, ecology and agriculture, underscores the importance of understanding the processes mediating adaptation to—and perhaps also by—WGD.

## Methods

### Plant material

We first located 89 *Cochlearia* populations throughout Europe and collected population samplings of plants from each, aiming for at least 10 plants per population, with each sampled plant a minimum of 2 meters from any other. Of these, we selected 33 geographically and ecotypically diverse representative populations for population resequencing, including the outgroup *Ionopsidium* (Dataset S2 Sample Metrics). An average of 4 individuals per population were sequenced. A total of 149 individuals were initially sequenced, which was narrowed down by a cutoff of minimum average depth of 4x, leaving 113 individuals from 33 populations in the final analysed dataset, including the outgroup *Ionopsidium*.

### Ploidy Determination by flow cytometry

DNA content and ploidy were inferred for populations using flow cytometry (Dataset S1). Approximately 1 square cm of leaf material was diced alongside an internal reference using razor blades in 1 ml ice cold extraction buffer (either 45 mM MgCl_2_, 30 mM sodium citrate, 20mM MOPS, 1% Triton-100, pH 7 with NaOH for relative staining or 0.1 M citric acid, 0.5% Tween 20 for absolute measurements). The resultant slurry was then filtered through a 40-μm nylon mesh before the nuclei were stained with the addition of 1 ml staining buffer (either CyStain UV precise P [Sysmex, Fluorescence emission: 435nm to 500nm] for relative ploidy, or Otto 2 buffer [0.4 M Na_2_HPO_4_·12H_2_O, Propidium iodide 50 μgmL−1, RNase 50 μgmL−1], for absolute DNA content). After 1 minute of incubation at room temperature the sample was run for 5,000 particles on either a Partec PA II flow cytometer or a BD FACS Melody. Histograms were evaluated using FlowJo software version 10.6.1.

### HEI10 immunostaining

Pachytene chromosome spreads were prepared from fixed anthers, as described previously^117^. Immunostaining was conducted using two primary antibodies: anti-AtZYP1C rat, 1:500^118^ and anti-HvHEI10 rabbit, 1:250^119^, followed by two secondary antibodies: goat anti-rat Alexa Fluor® 594 (Invitrogen) and goat anti-rabbit Alexa Fluor® 488 (Invitrogen). A Nikon Eclipse Ci fluorescence microscope equipped with NIS elements software was used to capture and quantify images. HEI10 foci were counted at pachytene using NIS software and significance established using the Mann-Whitney U test (Minitab v 18.1.0.0).

### Fluorescence *in situ* hybridization

Mitotic chromosome spreads from fixed root tips were prepared as described previously^120^. The *Arabidopsis*-type telomere repeat (TTTAGGG)_n_ was prepared according to ^121^. The *Cochlearia*-specific 102-bp (GTTAGATGTTTCATAAGTTCGTCAA ACTTGTACAAAGCTCATTGAGACACTTATAAGCACTCATGTTGCATGAAACTTGGTTTAGAGTCCTAGAAACGCGTT) tandem repeat was designed and prepared based on Mandáková et al. (2013) and used for identification of centromeres. The DNA probes were labeled by nick translation with biotin-dUTP and digoxigenin-dUTP according to ^122^, pooled and precipitated by adding 1/10 volume of 3 M sodium acetate, pH 5.2, and 2.5 volumes of ice-cold 96% ethanol and kept at −20°C for 30 min. The pellet was then centrifuged at 13,000 g at 4°C for 30 min. The pellet was resuspended in 20 µl of the hybridization mix (50% formamide and 10% dextran sulfate in 2×SSC) per slide. 20 µl of the probe was pipetted onto a chromosome-containing slide. The cover slips were framed with rubber cement. The probe and chromosomes were denatured together on a hot plate at 80°C for 2 min and incubated in a moist chamber at 37°C overnight. Post-hybridization washing was performed according to ^122^. After immunodetection, chromosomes were counterstained with 4’, 6-diamidino-2-phenylindole (DAPI, 2 µg/ml) in Vectashield (Vector Laboratories). The preparations were photographed using a Zeiss Axioimager Z2 epifluorescence microscope with a CoolCube camera (MetaSystems). The three monochromatic images were pseudocolored, merged and cropped using Photoshop CS (Adobe Systems) and Image J (National Institutes of Health) softwares.

### Reference Genome Assembly and Alignment

We generated a long read-based *de novo* genome assembly using Oxford Nanopore and Hi-C approaches, below.

### High Molecular Weight DNA isolation and Oxford Nanopore sequencing

A total of 0.4 g *Cochlearia excelsa* leaf material from one individual plant was ground using liquid nitrogen before the addition of 10 ml of CTAB DNA extraction buffer (100 mM Tris-HCl, 2% CTAB, 1.4 M NaCl, 20 mM EDTA, and 0.004 mg/ml Proteinase K). The mixture was incubated at 55°C for 1 hour then cooled on ice before the addition of 5 ml Chloroform. This was then centrifuged at 3000 rpm for 30 minutes and the upper phase taken, this was added to 1X volume of phenol:chloroform:isoamyl-alcohol and spun for 30 minutes at 3000 rpm. Again, the upper phase was taken and mixed with a 10% volume of 3M NaOAc and 2.5X volume of 100% ethanol at 4 °C. This was incubated on ice for 30 minutes before being centrifuged for 30 minutes at 3000 rpm and 4 °C. Three times the pellet was washed in 4ml 70% ethanol at 4 °C before being centrifuged again for 10 minutes at 3000 rpm and 4°C. The pellet was then air dried and resuspend in 300 µl nuclease-free water containing 0.0036 mg/ml RNase A. The quantity and quality of high molecular weight DNA was checked on a Qubit Fluorometer 2.0 (Invitrogen) using the Qubit dsDNA HS Assay kit. Fragment sizes were assessed using a Q-card (OpGen Argus) and the Genomic DNA Tapestation assay (Agilent). Removal of short DNA fragments and final purification to HMW DNA was performed with the Circulomics Short Read Eliminator XS kit.

Long read libraries were prepared using the Genomic DNA by Ligation kit (SQK-LSK109; Oxford Nanopore Technologies) following the manufacturer’s procedure. Libraries were then loaded onto a R9.4.1 PromethION Flow Cell (Oxford Nanopore Technologies) and run on a PromethION Beta sequencer. Due to the rapid accumulation of blocked flow cell pores or due to apparent read length anomalies on some *Cochlearia* runs, flow cells used in runs were treated with a nuclease flush to digest blocking DNA fragments before loading with fresh library according to the Oxford Nanopore Technologies Nuclease Flush protocol, version NFL_9076_v109_revD_08Oct2018.

### Genome size estimation and computational ploidy inference

We used KMC^123^ to create a k-mer frequency spectrum (Kmer length=21) of trimmed Illumina reads. We then used GenomeScope 2.0 (parameters: -k 21 -m 61) and Smudgeplot^124^ to estimate genome size and heterozygosity from k-mer spectra.

### Data processing and assembly

Fast5 sequences produced by PromethION sequencing were base called using the Guppy 6 high accuracy base calling model (dna_r9.4.1_450bps_hac.cfg) and the resulting fastq files were quality filtered by the base caller. A total of 17.2 GB base called data were generated for the primary assembly, resulting in 60x expected coverage. Primary assembly was performed in Flye^32^ and Necat^33^. The contigs were polished to improve the single-base accuracy in a single round of polishing in Medaka^34^ and Pilon^35^.

### Pseudomolecule construction by Hi-C, assembly cleanup, and polishing

To scaffold the assembled contigs into pseudomolecules, we performed chromosome conformation capture using HiC. Leaves from a single plant were snap-frozen in liquid N and ground to a fine powder using mortar and pestle. The sample was then homogenised, cross-linked and shipped to Phase Genomics (Seattle, USA), who prepared and sequenced an in vivo Hi-C library. After filtering low-quality reads with Trimmomatic^125^, we aligned the Hi-C reads against the contig-level assembly using bwa-mem^126^ (settings -5 -S -P) and removed PCR duplicated using Picard Tools (https://broadinstitute.github.io/picard/). We used 3D-DNA^127^ to conduct the initial scaffolding, followed by a manual curation in Juicebox^128^. After manually assigning chromosome boundaries, we searched for centromeric and telomeric repeats to orient the chromosome arms and to assess the completeness of the assembled pseudomolecules. To identify the centromeric repeat motif in *C. excelsa*, we used the RepeatExplorer^129^ pipeline to search for repetitive elements from short-read sequence data originating from the reference individual. RepeatExplorer discovered a highly abundant 102 nucleotide repeat element (comprising 21% of the short-read sequence), which we confirmed as the centromeric repeat motif by fluorescence in situ hybridisation. Using BLAST, we localised the centromeric and telomeric (TTTAGGG) repeats and used them to orient the chromosome arms. We performed a final assembly cleanup in Blobtools^36^ (Fig.S1). Gene space completeness was assessed using BUSCO version 3.0.2)^37^.

### Assembly annotation and RNA-seq

Prior to gene annotation, we identified and masked transposable element (TE) sequences from the genome assembly. To do so, we used the EDTA pipeline^130^, which combines multiple methods to comprehensively identify both retrotransposons and DNA transposons. After running EDTA on our chromosome-level genome assembly, we performed BLAST queries against a curated protein database from Swiss-Prot to remove putative gene sequences from the TE library and masked the remaining sequences from the assembly using RepeatMasker (https://www.repeatmasker.org).

We then used the BRAKER2^38^ pipeline to conduct gene annotatation on the TE-masked genome assembly. Evidence types included RNAseq data from the identical *C. excelsa* line and protein data from related species. RNA-seq was generated from bud, stem and leaf tissue. Total RNA was extracted from each tissue using the Qiagen RNeasy Extraction Kit. Stranded RNA libraries with polyA were constructed Using NEB Next Ultra II Directional RNA Library Prep Kit for Illumina and then evaluated by qPCR, TapeStation and Qubit at the DeepSeq facility (Nottingham, UK) before being sequenced at PE 150 at Novogene, inc (Cambridge, UK). We mapped the RNA-seq reads of each tissue to our reference genome using STAR^131^ with default parameters (-twopassMode Basic) before running BRAKER2. Running BRAKER2 without UTR prediction generated more gene models and much better BUSCO metrics than with UTR prediction (97.8% [raw, pre-Blobtools trimmed] complete BUSCOs without UTR prediction vs 91.7% with UTR prediction), so for the final annotation we used the more complete set and ran BRAKER2 without UTR prediction.

### Population Resequencing and Analysis

#### Library preparation and sequencing

DNA was prepared using the commercially available DNeasy Plant Mini Kit from Qiagen. DNA libraries were made using TruSeq DNA PCR-free Library kit from Illumina as per the manufacturer’s instructions and were multiplexed based on concentrations measured with a Qubit Flourometer 2.0 (Invitrogen) using the Qubit dsDNA HS Assay kit. Sequencing was carried out on either NextSeq 550 (Illumina) in house (4 runs) or sent to Novogene for Illumina Hiseq X, PE150 sequencing (2 runs).

#### Data preparation, alignment, and genotyping

Reads were quality trimmed with Trimmomatic 0.39^125^ (PE -phred33 LEADING:10 TRAILING:10 SLIDINGWINDOW:4:15 MINLEN:50) and then aligned to the *C. excelsa* reference using bwa-mem^132^ and further processed with samtools^133^. Duplicate reads were removed and read group IDs added to the bam files using Picard (version 1.134). Indels were realigned with GATK (version 4.2.3.0)^134^. Samples were first genotyped individually with “HaplotypeCaller” (--emit-ref-confidence BP_RESOLUTION --min-base-quality-score 25 --minimum-mapping-quality 25) and were then genotyped jointly using “GenotypeGVCFs” in GATK (version 4.2.3.0). The resulting VCF files were then filtered for biallelic sites and mapping quality (QD < 2.0, FS > 60.0, MQ < 40.0, MQRankSum < −12.5, ReadPosRankSum < −8.0, HaplotypeScore < 13.0). The VCF was then filtered by depth. Based on this distribution a depth cutoff of 4,322 was applied to the VCF containing the dataset and this was then used as a mask for the final VCF containing all individuals.

#### Demographic analysis

We first inferred genetic relationships between individuals using principal component analysis (PCA). Following ^135^, we estimated a matrix of genetic covariances between each pair of individuals. For two individuals, 𝑖 and 𝑗, covariance (𝐶) was calculated as:

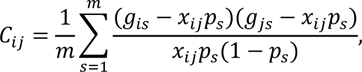

where 𝑚 is the number of variable sites, 𝑔_!#_ is the genotype of individual 𝑖 in site 𝑠, 𝑥 is the average ploidy level of the two individuals, and 𝑝 is the alternate allele frequency. PCA was performed on the matrix using the R function *prcomp*, setting scaling to TRUE, and first two axes of the rotated data extracted for plotting. For fastSTRUCTURE we followed ^136^ by randomly subsampling two alleles from tetraploid and hexaploid populations using a custom script. We have previously demonstrated that results generated in this way are directly comparable to results generated with the full dataset in STRUCTURE^136^. We calculated Nei’s distances among all individuals in StAMPP^137^ and visualized these using SplitsTree^39^. Linkage disequilibrium was estimated using ldsep^138^. To avoid biasing the estimates with unequal sample sizes, we chose 39 diploids and tetraploids for the analysis. To reduce computation time, the analysis was performed on 4-fold sites from a single chromosome (chromosome 1). To visualize the decay of LD as a function of physical distance, we calculated average *r*^2^ in 10 bp non-overlapping windows and fit a loess curve on the binned data.

#### Window-based scan for selective sweep signatures

We performed a window-based divergence scan for selection consisting of 1 kb windows that contained at least 15 SNPs. The data were filtered as described above and in addition was filtered for no more than 20% missing data and a depth of >= 8x. We calculated the following metrics: Rho^45^, Nei’s F_ST_ ^47^, Weir-Cochran’s F_ST_^48^, Hudson’s F_ST_^46^, Dxy^44^, number of fixed differences and average groupwise allele frequency difference (AFD). To determine the best metric to detect localised peaks of divergence we performed a quantitative analysis of AFD plot quality for all 1% outliers of each metric. Each window was given a score of 0-4, with 0 being the lowest quality and 4 the highest. Scores were based on two qualities: peak height and peak specificity. For peak height one point was awarded if the window contained one SNP of AFD between 0.5 and 0.7, and two points were awarded for any SNP of AFD > 0.7. Likewise, for peak specificity two points were awarded for an AFD peak that was restricted to a single gene and one point was awarded for a peak that was restricted to 2-3 genes. Compared to all other single 1% outlier lists and all permutations of overlapped 1% outlier lists, the top 1% outliers from Hudson’s F_ST_ performed most favorably as it maximized the number of ‘4’ and ‘3’ scores while minimizing the number of ‘1’ and ‘0’ scores. Finally, we masked from downstream analysis a region of uniformly high differentiation marking a suspected inversion at scaffold 6 (between 5,890,246 and 6,137,362 bp).

#### MAV analysis

Following ^13^, we performed a FineMAV^50^-like analysis on all biallelic, non-synonymous SNPs passing the same filters as the window-based selection scan. SNPs were assigned a Grantham score according to the amino acid change and this was scaled by the AFD between ploidies. SNPs were first filtered for a minimum AFD of 0.25. The top 1% outliers of all these MAV-SNPs were then overlapped with the genes in our 1% F_ST_ outlier windows to give a refined list of candidate genes that contain potentially functionally significant non-synonymous mutations at high AFD between cytotypes. The code outlining this can be found at https://github.com/paajanen/meiosis_protein_evolution/tree/master/FAAD.

#### Orthogrouping and Reciprocal Best Blast Hits

We performed an orthogroup analysis using Orthofinder version 2.5.5^110^. to infer orthologous groups (OGs) from four species (*C. amara, A. lyrata, A. thaliana, C. excelsa*). A total of 25,199 OGs were found. Best reciprocal blast hits (RBHs) for *Cochlearia* and *A. thaliana* genes were found using BLAST version 2.9.0. *Cochlearia* genes were then assigned an *A. thaliana* gene ID for GO enrichment analysis in one of five ways. First if the genes’ OG contained only one *A. thaliana* gene ID, that gene ID was used. If the OG contained more than one *A. thaliana* gene ID then the RBH was taken. If there was no RBH then the OG gene with the lowest E-value in a BLAST versus the TAIR10 database was taken. If no OG contained the *Cochlearia* gene then the RBH was taken. Finally, if there was no OG or RBH then the gene with the lowest E-value in a BLAST versus the TAIR10 database was taken. BLASTs were performed using the TAIR10.1 genome with data generated on 2023-01-02.

#### GO Enrichment Analysis

To infer functions significantly associated with directional selection following WGD, we performed gene ontology enrichment of candidate genes in the R package TopGO v.2.52^139^, using our *Cochlearia* universe set. We tested for overrepresented Gene Ontology (GO) terms within the three domains Biological Process (BP), Cellular Component (CC) and Molecular Function (MF) using Fisher’s exact test with conservative ‘elim’ method, which tests for enrichment of terms from the bottom of the hierarchy to the top and discards any genes that are significantly enriched in a descendant GO term. We used a significance cut-off of 0.05.

#### Generation of Consensus Sequences

Consensus sequences were generated for proteins of interest so that they could be closely inspected via MSAs and 3D protein structure prediction. Genomic regions were selected for either all diploids or all tetraploids present in the selection scan with GATK SelectVariants, while simultaneously being filtered for biallelic SNPs, “--max-nocall-fraction 0.2” and “-select ‘AF > 0.5’”. A consensus sequence was generated for exons by combining samtools faidx and bcftools consensus. Finally, a VCF containing only non-biallelic variation was manually inspected and any multiallelic variants at AF>0.5 and max-nocall-fraction<0.2 manually incorporated into the consensus.

#### Multiple Sequence Alignments (MSAs)

We generated multiple sequence alignments using Clustal-Omega^140^ in combination with amino acid sequences from the GenBank database. Sequences were selected either because the genes/proteins were well studied in other organisms or to give a phylogenetically broad coverage. Alignments were manually refined and visualised in JalView^141^.

#### Protein modeling

Protein homology models were created using AlphaFold^142^ version 2.1 on the Czech national HPC MetaCentrum. The full database was uses with a model preset of monomer and a maximum template data of 2020-05-14. Structures were visualised and images generated in the PyMOL Molecular Graphics System (Version 2.0 Schrödinger, LLC).

## Data Availability

All sequence data for this study have been deposited in the European Nucleotide Archive (ENA; https://www.ebi.ac.uk/ena/browser/view/PRJEB66308), accession number PRJEB66308. The chromosome build genome assembly and annotation files are available at Data Dryad at https://doi.org/10.5061/dryad.ncjsxkt1s.

## Acknowledgements

We thank Kirsten Bomblies, Lara Hebberecht-Lopez, Mary Bray and Nigel Bray for assistance collecting plant material. We further acknowledge wet lab support from Anna Loreth (Heidelberg). This work was supported by the European Research Council (ERC) under the European Union’s Horizon 2020 research and innovation program [grant number ERC-StG 679056 HOTSPOT], via a grant to LY and under the Marie Skłodowska-Curie grant agreement No. 101022295 to TH. Nanopore sequencing was performed in the University of Nottingham DeepSeq sequencing facility. We are grateful for access to the University of Nottingham’s Augusta HPC service and the CESNET LM2015042 and the CERIT Scientific Cloud LM2015085, provided under the programme ‘Projects of Large Research, Development, and Innovations Infrastructures’. We thank local authorities for issuing permission to collect samples (Slovak Ministry for Environment, permission No 062-219/18). The article is available in gold open access.

## Author Contributions

LY and SMB conceived the study. LY, SMB, MK, PM, and JK performed collections. SMB, SB, SDD, T.Mandáková, CM, LC, MZ, and SF performed laboratory experiments. SMB, TH, SDD, CM, T.Mathers, T.Mandáková, EW, MB, SB, SF, PP, MK and LY performed analyses. LY and SMB wrote the manuscript. All authors approved the final manuscript.

