## Supplemental Text Figures and Tables for "Kinetochore and ionomic adaptation to whole genome duplication"

### Supplementary Text

#### Supplementary Text 1: Salinity tolerance experiments.

Seeds were sowed in petri plates filled with wet sand for a week, then transferred to pot trays filled with John Innes mix (No.1). Each tray contained 60 plants. A total of 1,036 plants were planted. The experiment was conducted in complete randomized design with as many replications as possible per maternal seed line (average 13.05) and plants were grown in a glasshouse at ambient UK summer temperature (~25C day, ~10C night). After two weeks from germination, trays were bottom-watered twice weekly with watering solution containing either 10 mM MES pH 7 and NaCl for the treatment or just 10 mM MES for the control. Salt concentrations began at 50 mM NaCl for two weeks, then were increased to 300 mM for two weeks, and were finally increased to 600 mM for 4 weeks. During the final week, leaves were sprayed with 600 mM NaCl watering solution twice. After 8 total weeks of treatment, a modified Standard Evaluation System (SES) Score was used to evaluate symptoms of salt stress[1]. Three individuals independently took measurements for each plant and the final score was given as the average of the three. At the same time point, two leaves of each plant were harvested, dried at 60 ºC for two days and stored for ICP analysis. The diploids were significantly more salt tolerant than the tetraploids (W = 29568, p-value = 2.178e-05, Wilcoxon rank sum test with continuity correction performed in R).

#### Supplementary Text 2: Stomatal conductance assessment, net photosynthesis and drought tolerance experiments.

To test for differences in physiological responses to drought stress we cultivated twenty different populations of *Cochlearia* side by side in a controlled environment. Seeds were germinated in wet sand in a growth camber with temperature and humidity control before being transplanted into PET-based potting soil and placed in a growth room (16h light from 6:00 to 20:00, temperature day: 22°C and night: 18°C). The watering regime for control and drought stressed plants was as follows: For the first two weeks after transplanting, they were watered 1000 ml $H_2O$ per tray every 2nd day. From the 3rd week onwards a drought stress was imposed. Control plants were still watered with 1000 ml $H_2O$ per tray every 2nd day. Drought stress was generated by lowering watering/soil moisture week by week as follows: week 3 - 1000 ml $H_2O$ per tray every 3rd day, week 4 – 1000 ml $H_2O$ per tray every 4th day, week 5 – 1000 ml $H_2O$ per tray every 5th day, week 6 – 1000 ml $H_2O$ per tray every 6th day. Plants were assessed at the end of week 6. Wet and dry weight of the shoot as well as photosynthesis parameters were captured. Photosynthetic parameters were assessed using the LI-6400XT Portable Photosynthesis System (LI-COR, Lincoln, NE, United States). All measurements were taken between 9:30 and 11:30 using light adapted plants. Photosynthetic parameters were measured on the largest leaf from both control and drought treatment groups and all measurements were conducted at the same time of the day (9:30am-11:30am). The leaf was placed into a custom-made single leaf chamber (chamber: polyoxymethylene, lid: poly (methyl methacrylate)), openings sealed with sponge rubber, which was connected to the LI-COR. Photosynthetic parameters were recorded after the reads were stable. We analysed 9 diploid populations and 11 tetraploid populations. We observed an increased net photosynthetic rate in tetraploids in both control and drought conditions. We note large population-specific variation in both cytotypes.

Figure 2E and F show stomatal conductance (SC) of diploid and tetraploid populations under drought stress relative to well-watered conditions (relative SC). Averages, standard deviation and confidence interval for relative stomatal conductance and net photosynthesis can be found in Tables S1-S3. Tetraploids show higher SC under drought stress. As the linear model in Table S4 shows, ploidy significantly affects the phenotype.

Data were analysed using R Version 3.6.0 (2019-04-26) with RStudio Version 1.2.1335. Data were plotted using the ggplot2 package version 3.2.1. The lsmeans package version 2.30-0 was used to fit a linear model to the data while the package bestNormalize version 1.4.3 was used for data normalization and estimation of normality. The package sjPlot version 2.8.2 was used to generate a summary table for the linear model fitted to the data. The package flextable version 0.5.9 was used to generate summary tables containing averages, sd and ci intervals.

1. Gregoria, G. B., Senadhira, D. & Mendoza, R. D. *Screening rice for salinity tolerance*. https://econpapers.repec.org/RePEc:ags:irridp:287589 (1997).

### Supplemental Figures

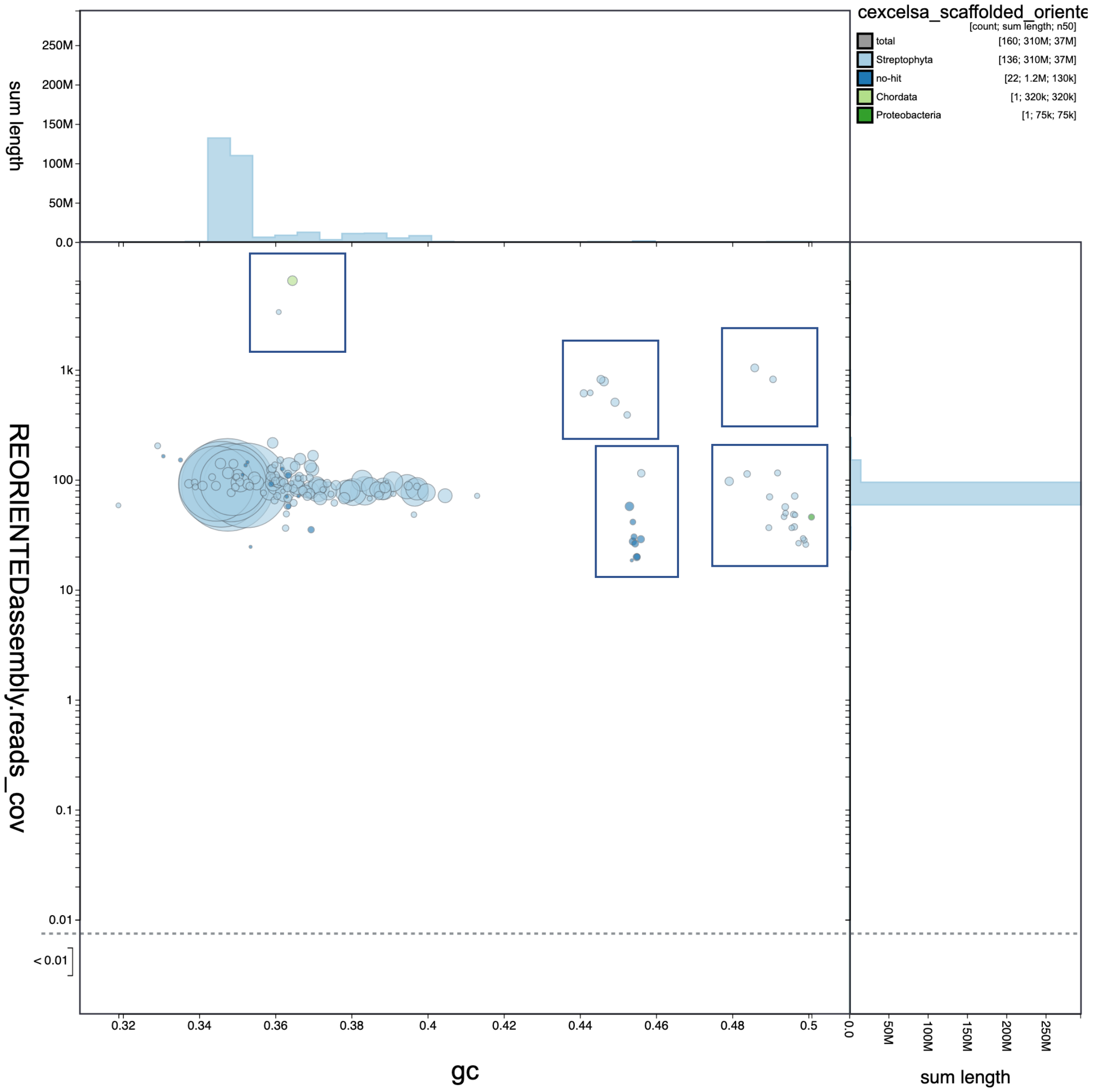


**Figure S1. *Cochlearia excelsa* genome assembly cleanup.** Final filtering of contaminant contigs with Blobtools. Rectangles indicate contigs flagged as contamination or organellar based on GC content (x-axis), outlier read depth (y-axis), and homology in the UniProt database (colours, in key insert top right; see methods).

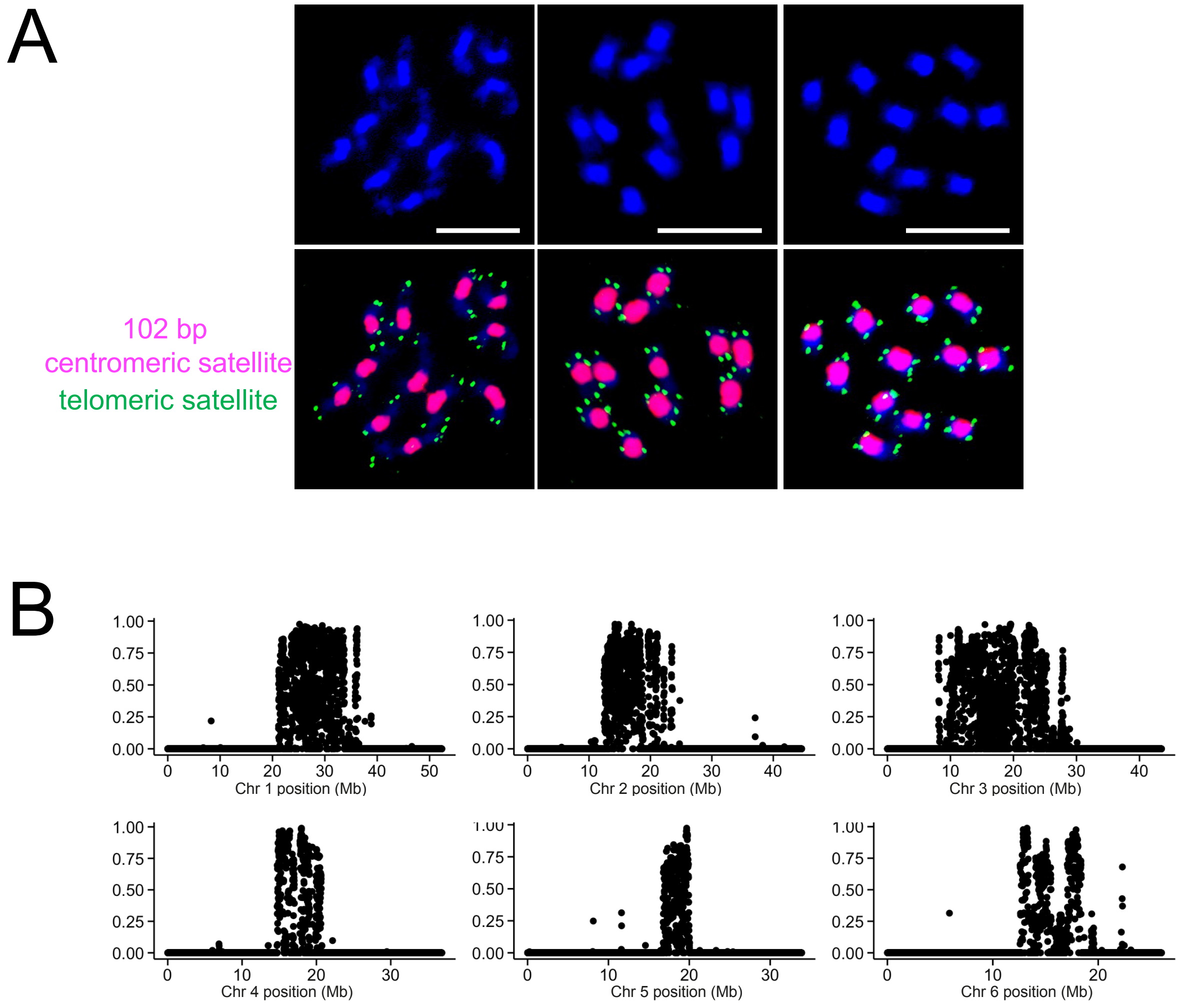


**Figure S2. FISH and repeat mapping in *Cochlearia excelsa* (2n = 12). A)** Cytology shows DAPI-stained chromosomes (2n = 12) with heterochromatic pericentromeres (top) and 102 bp satellite (pink) and *Arabidopsis*-type telomeric satellite (green) probes hybridizing to all (peri)centromeres and telomeres, respectively (bottom). Scale bar, 10 μm; **B)** Mapping of simple 102 bp repeat, comprising 21% of genomic sequence.

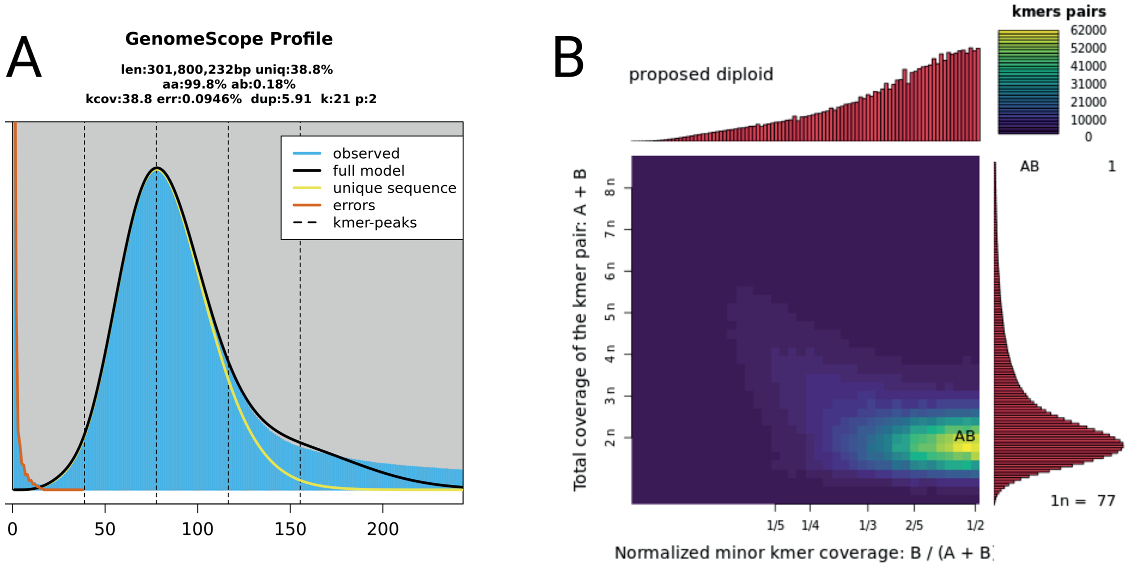


**Figure S3. Genome size and ploidy estimates of *Cochlearia excelsa* chromosome build genome.** (A) Haploid genome size of 302 mb and very low heterozygosity using Genomescope and (B) ploidy confirmation by Smudgeplot

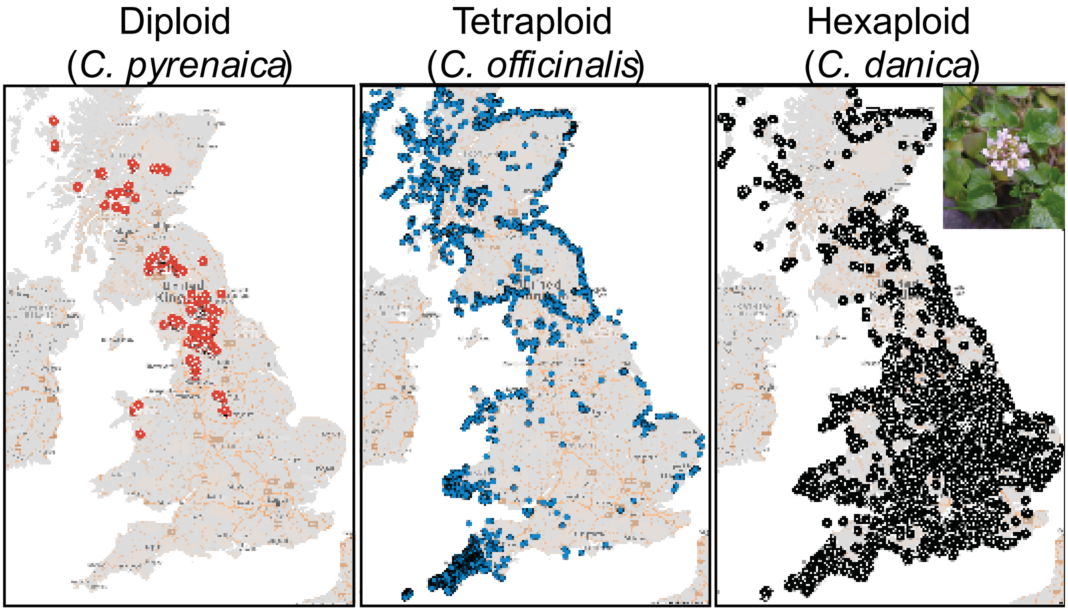


**Figure S4. Contrasting distributions of major *Cochlearia* cytotypes.** Left: Diploids deeply inland in nonsaline habitats; Centre; tetraploids inhabiting coastal regions; Right: Cochlearia danica, the hexaploid, exclusively coastal until the 1970's, has explosively invaded salted motorways since the practice of salting the roads began at that time. Data: complete download of reported botanical observations from the records of the Botanical Society of Britain & Ireland (1895-2021).

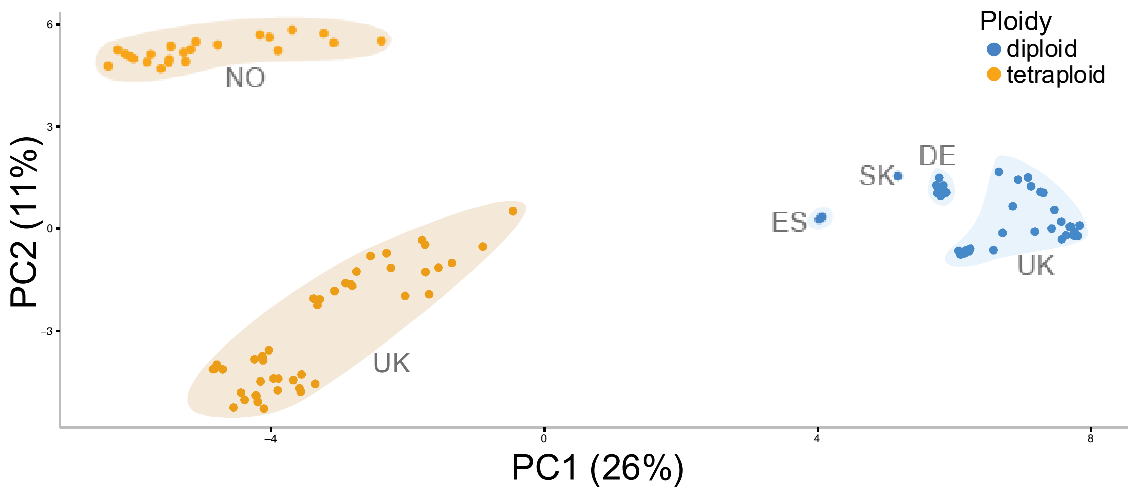


**Figure S5. Principal component analysis of Cochlearia diploids (blue) and autotetraploids (orange).** PC1 explains 26% of the variation and discriminates populations based on ploidy, with autotetraploids (orange) on the left. PC2 explains only 11% of the variation and separates the Norwegian tetraploids from the UK tetraploids. Axes are scaled to contribution of each of the first principal components to the overall variation. NO=Norway, UK=United Kingdom, ES=Spain, SK=Slovakia, DE=Germany.

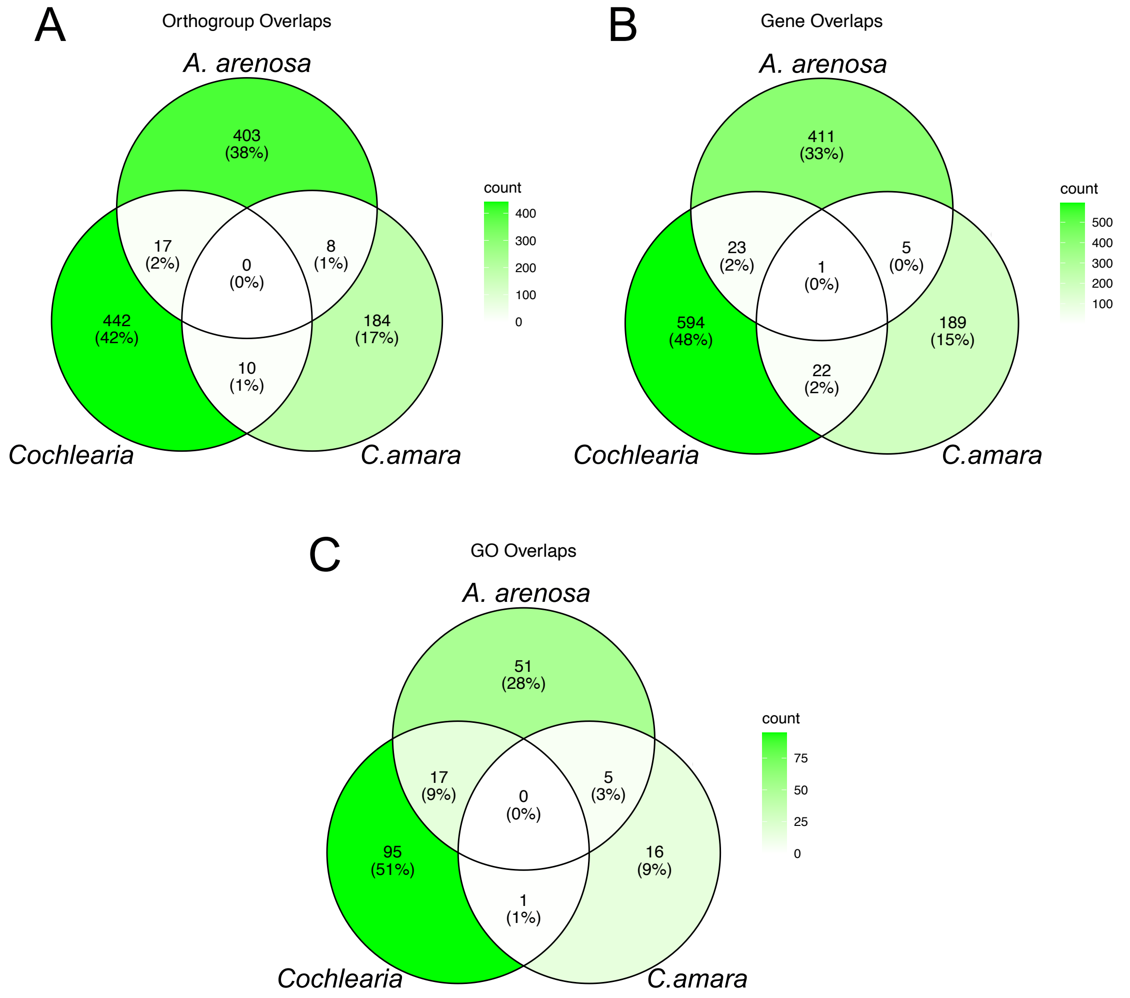


**Figure S6. Process-level, but not gene-level convergence.** Overlaps of *Cochlearia, A. arenosa,* and *C. amara* selective sweep candidates by the same metric, Hudson's Fst. A modest degree of process-level convergence (panel A) is evident between *Cochlearia* and *A. arenosa*. A lack of enrichment for gene-level convergence is seen either by the orthogrouping (Panel B) or nearest homolog (Panel C) approaches.

### Supplemental Tables

**Table S1.** Phenotype scores for *Cochlearia* plants following salt stress treatment.

| Maternal Line | Scores | Average Score | Number of Samples | Standard Deviation | Ploidy |
|---|---|---|---|---|---|
| **CHA_001** | 7, 5, 6, 7, 5, 7, 7, 4, 4, 3, 5, 8, 0, 0, 3 | 4.73 | 15 | 2.46 | 2x |
| **CHA_010** | 5, 7, 5, 6, 4, 2, 3 | 4.57 | 7 | 1.72 | 2x |
| **CHA_011** | 7, 4, 0, 4, 6, 6, 8, 7, 2, 2, 4, 5, 3 | 4.46 | 13 | 2.33 | 2x |
| **CHA_002** | 4, 6, 6, 7, 5, 5, 6, 6 | 5.63 | 8 | 0.92 | 2x |
| **CHA_008** | 7, 7, 8, 8, 8, 8, 7, 7, 9, 9, 7, 7, 9, 6, 9, 8, 6, 8, 8 | 7.68 | 19 | 0.95 | 2x |
| **CWY_001** | 1 | 1.00 | 1 | N/A | 4x |
| **CWY_013** | 3, 3, 3, 2, 2, 3, 2, 2, 5, 5, 5 | 3.18 | 11 | 1.25 | 4x |
| **CWY_002** | 2 | 2.00 | 1 | N/A | 4x |
| **CWY_003** | 3, 4, 5, 5, 2, 4, 3, 4, 4, 2, 3, 2, 1 | 3.23 | 13 | 1.24 | 4x |
| **CWY_005** | 4, 3, 1, 2, 3, 2, 2, 3, 1, 1, 1 | 2.09 | 11 | 1.04 | 4x |
| **LAB_002** | 5, 7, 7 | 6.33 | 3 | 1.15 | 2x |
| **LAB_003** | 6, 4, 8, 6, 5, 5, 6, 8, 6, 5, 5, 4, 6, 2, 7, 4, 5, 2, 4, 7 | 5.25 | 20 | 1.65 | 2x |
| **LAB_007** | 5, 6, 5, 7, 4, 6, 3, 6, 5, 3, 4, 4, 5, 5, 6, 6 | 5.00 | 16 | 1.15 | 2x |
| **LAB_008** | 8, 6, 6, 7, 7, 7, 8, 8, 5, 7, 6, 8, 7, 6, 7, 7, 5, 7, 7, 7 | 6.80 | 20 | 0.89 | 2x |
| **LAB_009** | 6, 5, 4, 5, 5, 5, 6, 7, 8, 5, 6, 5, 6, 5, 4, 5 | 5.44 | 16 | 1.03 | 2x |
| **LAM_017** | 8, 6 | 7.00 | 2 | 1.41 | 2x |
| **MAL_002** | 6, 7, 5, 6, 6, 7, 8, 6, 7, 5, 2, 0, 4, 6, 6, 7 | 5.50 | 16 | 2.03 | 2x |
| **MAL_003** | 7, 8, 8, 5, 5, 5, 8, 8, 8, 5, 6, 9, 5 | 6.69 | 13 | 1.55 | 2x |
| **MAL_004** | 7, 6, 6, 6, 4, 4, 6, 7, 6, 7, 7 | 6.00 | 11 | 1.10 | 2x |
| **MAL_007** | 6, 5, 7, 6, 8, 6, 3, 4, 6, 6, 4, 6, 1, 3, 5 | 5.07 | 15 | 1.79 | 2x |
| **NEN_002** | 1, 3, 2, 2, 3, 3, 3, 3, 1, 2, 1, 1, 2, 3, 3, 2, 3, 1, 2, 1 | 2.10 | 20 | 0.85 | 2x |
| **NEN_003** | 7, 8, 8, 6, 6, 6, 7, 5, 8, 5, 4, 6, 8, 7, 0, 3, 7, 8, 5, 6 | 6.00 | 20 | 2.00 | 2x |
| **NEN_004** | 2, 5, 7, 5, 4, 6, 7, 5, 5, 8, 8, 6, 6 | 5.69 | 13 | 1.65 | 2x |
| **NEN_005** | 9, 6, 3, 7, 4, 6, 5, 7, 5, 8, 9, 6, 7, 3, 0, 4 | 5.56 | 16 | 2.39 | 2x |
| **NEN_006** | 4, 6, 5, 6 | 5.25 | 4 | 0.96 | 2x |
| **SCU_012** | 4, 3, 5, 4, 6, 4, 5, 4, 5, 5, 6, 7, 5, 5, 5, 5, 0, 5, 4, 7 | 4.70 | 20 | 1.49 | 4x |
| **SCU_020** | 6, 7, 7, 7, 7, 7, 9, 7, 8, 6, 9, 10, 8, 9, 8, 8, 7, 9, 9 | 7.79 | 19 | 1.13 | 4x |
| **SCU_003** | 4, 4, 3, 1 | 3.00 | 4 | 1.41 | 4x |
| **SCU_009** | 3, 2, 3, 5 | 3.25 | 4 | 1.26 | 4x |
| **TMN_001** | 3, 3, 4, 6, 3, 3, 5, 5, 4, 3, 2, 3, 0, 2, 3, 3, 2, 3, 4 | 3.21 | 19 | 1.32 | 4x |
| **TMN_003** | 4, 5, 5, 5, 3, 3, 3, 2, 6, 3, 3, 5, 4, 4, 5, 4, 5, 7, 5, 4 | 4.25 | 20 | 1.21 | 4x |
| **TMN_005** | 1, 5, 2, 1 | 2.25 | 4 | 1.89 | 4x |
| **TMN_006** | 5, 4 | 4.50 | 2 | 0.71 | 4x |
| **TYN_0H1** | 1, 4, 8, 5, 4, 2, 2, 2, 3, 2, 2, 1, 1, 3, 1, 2, 1, 2, 1, 1 | 2.40 | 20 | 1.76 | 2x |
| **TYN_0H4** | 2, 3, 1, 3, 2, 1, 3, 1, 0, 1, 2, 4, 2, 1, 3, 2, 1, 2, 2, 0 | 1.80 | 20 | 1.06 | 2x |
| **TYN_0H5** | 1, 3, 9, 7, 9, 8, 4, 0, 7, 8, 8, 8, 7, 6, 8, 9, 5, 5, 6, 5 | 6.15 | 20 | 2.58 | 2x |
| **TYN_0H7** | 0, 5, 6, 6, 7, 5, 5, 4, 3, 0, 4, 6, 3, 2, 4, 5, 4, 2, 5, 3 | 3.95 | 20 | 1.90 | 2x |
| **TYN_0H8** | 7, 6, 3, 4, 3, 3, 4, 5, 6, 6, 4, 5, 4, 3, 0, 1, 6, 5, 6, 6 | 4.35 | 20 | 1.81 | 2x |
| **All diploids** | N/A | 4.98 | 367 | 1829.00 | 2x |
| **All tetraploids** | N/A | 4.14 | 129 | 534.00 | 4x |

**Table S2**. Net Photosynthetic rates under drought relative to control conditions.

| relative net Photosynthesis | | | | | | | |
|---|---|---|---|---|---|---|---|
| **Species** | **Treatment** | **Ploidy** | **N** | **Average** | **sd** | **se** | **ci** |
| ELI21 | Control | 4x | 4 | **1.00** | 0.12 | 0.06 | 0.19 |
| ELI21 | Drought | 4x | 5 | **0.91** | 0.10 | 0.04 | 0.12 |
| ELI23 | Control | 4x | 3 | **1.00** | 0.16 | 0.09 | 0.40 |
| ELI23 | Drought | 4x | 4 | **0.78** | 0.04 | 0.02 | 0.06 |
| ELI25 | Control | 4x | 5 | **1.00** | 0.22 | 0.10 | 0.27 |
| ELI25 | Drought | 4x | 3 | **0.99** | 0.09 | 0.05 | 0.22 |
| ERS21+22 | Control | 4x | 4 | **1.00** | 0.05 | 0.03 | 0.08 |
| ERS21+22 | Drought | 4x | 4 | **1.04** | 0.10 | 0.05 | 0.17 |
| ERS23 | Control | 4x | 3 | **1.00** | 0.19 | 0.11 | 0.48 |
| ERS23 | Drought | 4x | 4 | **0.66** | 0.04 | 0.02 | 0.07 |
| ERS26 | Control | 4x | 3 | **1.00** | 0.27 | 0.15 | 0.66 |
| ERS26 | Drought | 4x | 4 | **0.83** | 0.10 | 0.05 | 0.16 |
| LAB22 | Control | 2x | 5 | **1.00** | 0.25 | 0.11 | 0.31 |
| LAB22 | Drought | 2x | 5 | **0.77** | 0.11 | 0.05 | 0.14 |
| LAB23 | Control | 2x | 4 | **1.00** | 0.06 | 0.03 | 0.09 |
| LAB23 | Drought | 2x | 5 | **0.60** | 0.17 | 0.08 | 0.22 |
| Nent20 | Control | 2x | 3 | **1.00** | 0.03 | 0.02 | 0.08 |
| Nent20 | Drought | 2x | 7 | **0.54** | 0.14 | 0.05 | 0.13 |
| Nent23 | Control | 2x | 4 | **1.00** | 0.07 | 0.03 | 0.11 |
| Nent23 | Drought | 2x | 3 | **0.48** | 0.17 | 0.10 | 0.42 |
| Nent24 | Control | 2x | 4 | **1.00** | 0.32 | 0.16 | 0.51 |
| Nent24 | Drought | 2x | 6 | **0.86** | 0.12 | 0.05 | 0.12 |
| PIT_020 | Control | 2x | 13 | **1.00** | 0.14 | 0.04 | 0.09 |
| PIT_020 | Drought | 2x | 15 | **0.82** | 0.23 | 0.06 | 0.13 |
| PIT_022 | Control | 2x | 3 | **1.00** | 0.13 | 0.07 | 0.32 |
| PIT_022 | Drought | 2x | 3 | **0.58** | 0.08 | 0.05 | 0.20 |
| PIT_023 | Control | 2x | 3 | **1.00** | 0.03 | 0.02 | 0.08 |
| PIT_023 | Drought | 2x | 3 | **0.84** | 0.07 | 0.04 | 0.19 |
| PIT_025 | Control | 2x | 5 | **1.00** | 0.07 | 0.03 | 0.08 |
| PIT_025 | Drought | 2x | 5 | **0.62** | 0.04 | 0.02 | 0.05 |
| SKN20 | Control | 4x | 3 | **1.00** | 0.18 | 0.10 | 0.44 |
| SKN20 | Drought | 4x | 6 | **0.65** | 0.10 | 0.04 | 0.10 |
| SKN21-A | Control | 4x | 6 | **1.00** | 0.16 | 0.07 | 0.17 |
| SKN21-A | Drought | 4x | 6 | **0.87** | 0.20 | 0.08 | 0.21 |
| SKN21-B | Control | 4x | 3 | **1.00** | 0.14 | 0.08 | 0.34 |
| SKN21-B | Drought | 4x | 5 | **0.93** | 0.27 | 0.12 | 0.34 |
| SKN24 | Control | 4x | 5 | **1.00** | 0.12 | 0.06 | 0.15 |
| SKN24 | Drought | 4x | 5 | **0.82** | 0.16 | 0.07 | 0.20 |

**Note:** Averages, standard deviation and 95% confidence interval under control and drought stress are shown.

**Table S3.** Stomatal conductance under drought relative to control conditions.

| relative Stomata Conductance | | | | | | | | |
|---|---|---|---|---|---|---|---|---|
| **Species** | **Treatment** | **Ploidy** | **N** | **Average** | **sd** | **se** | **ci** | **rn** |
| ELI21 | Control | 4x | 4 | **1.00** | 0.15 | 0.08 | 0.24 | 1.00 |
| ELI21 | Drought | 4x | 5 | **0.70** | 0.20 | 0.09 | 0.24 | 0.70 |
| ELI23 | Control | 4x | 3 | **1.00** | 0.17 | 0.10 | 0.42 | 1.00 |
| ELI23 | Drought | 4x | 4 | **0.98** | 0.17 | 0.09 | 0.28 | 0.98 |
| ELI25 | Control | 4x | 5 | **1.00** | 0.20 | 0.09 | 0.24 | 1.00 |
| ELI25 | Drought | 4x | 3 | **1.06** | 0.95 | 0.55 | 2.37 | 1.06 |
| ERS21+22 | Control | 4x | 4 | **1.00** | 0.31 | 0.15 | 0.49 | 1.00 |
| ERS21+22 | Drought | 4x | 4 | **0.62** | 0.11 | 0.05 | 0.17 | 0.62 |
| ERS23 | Control | 4x | 3 | **1.00** | 0.39 | 0.22 | 0.97 | 1.00 |
| ERS23 | Drought | 4x | 4 | **1.01** | 0.50 | 0.25 | 0.79 | 1.01 |
| ERS26 | Control | 4x | 3 | **1.00** | 0.25 | 0.14 | 0.61 | 1.00 |
| ERS26 | Drought | 4x | 4 | **1.44** | 0.82 | 0.41 | 1.31 | 1.44 |
| LAB22 | Control | 2x | 5 | **1.00** | 0.48 | 0.21 | 0.60 | 1.00 |
| LAB22 | Drought | 2x | 5 | **0.53** | 0.24 | 0.11 | 0.30 | 0.53 |
| LAB23 | Control | 2x | 4 | **1.00** | 0.41 | 0.20 | 0.65 | 1.00 |
| LAB23 | Drought | 2x | 5 | **0.39** | 0.09 | 0.04 | 0.12 | 0.39 |
| Nent20 | Control | 2x | 3 | **1.00** | 0.00 | 0.00 | 0.01 | 1.00 |
| Nent20 | Drought | 2x | 7 | **0.52** | 0.18 | 0.07 | 0.16 | 0.52 |
| Nent23 | Control | 2x | 4 | **1.00** | 0.17 | 0.08 | 0.26 | 1.00 |
| Nent23 | Drought | 2x | 3 | **0.54** | 0.18 | 0.11 | 0.46 | 0.54 |
| Nent24 | Control | 2x | 4 | **1.00** | 0.27 | 0.13 | 0.43 | 1.00 |
| Nent24 | Drought | 2x | 6 | **0.71** | 0.18 | 0.07 | 0.19 | 0.71 |
| PIT_020 | Control | 2x | 13 | **1.00** | 0.23 | 0.06 | 0.14 | 1.00 |
| PIT_020 | Drought | 2x | 15 | **0.81** | 0.26 | 0.07 | 0.14 | 0.81 |
| PIT_022 | Control | 2x | 3 | **1.00** | 0.43 | 0.25 | 1.07 | 1.00 |
| PIT_022 | Drought | 2x | 3 | **1.18** | 0.05 | 0.03 | 0.12 | 1.18 |
| PIT_023 | Control | 2x | 3 | **1.00** | 0.37 | 0.21 | 0.92 | 1.00 |
| PIT_023 | Drought | 2x | 3 | **0.71** | 0.38 | 0.22 | 0.95 | 0.71 |
| PIT_025 | Control | 2x | 5 | **1.00** | 0.15 | 0.07 | 0.19 | 1.00 |
| PIT_025 | Drought | 2x | 5 | **0.28** | 0.04 | 0.02 | 0.05 | 0.28 |
| SKN20 | Control | 4x | 3 | **1.00** | 0.31 | 0.18 | 0.78 | 1.00 |
| SKN20 | Drought | 4x | 6 | **0.70** | 0.08 | 0.03 | 0.08 | 0.70 |
| SKN21-A | Control | 4x | 6 | **1.00** | 0.38 | 0.15 | 0.40 | 1.00 |
| SKN21-A | Drought | 4x | 6 | **0.84** | 0.36 | 0.15 | 0.37 | 0.84 |
| SKN21-B | Control | 4x | 3 | **1.00** | 0.16 | 0.09 | 0.39 | 1.00 |
| SKN21-B | Drought | 4x | 5 | **0.91** | 0.33 | 0.15 | 0.41 | 0.91 |
| SKN24 | Control | 4x | 5 | **1.00** | 0.12 | 0.05 | 0.14 | 1.00 |
| SKN24 | Drought | 4x | 5 | **0.57** | 0.13 | 0.06 | 0.16 | 0.57 |

**Note:** Averages, standard deviation and 95% confidence intervals under control and drought stress are shown.

**Table S4.** Stomatal conductance and photosynthesis model results.

| | Rel. Stomatal Conductance | | | rel. Photosynthesis | | |
|---|---|---|---|---|---|---|
| *Predictors* | *Estimates* | *CI* | *p* | *Estimates* | *CI* | *p* |
| (Intercept) | -0.29 | -0.55 – -0.03 | **0.030** | -0.31 | -0.57 – -0.05 | **0.021** |
| Ploidy [4x] | 0.62 | 0.24 – 1.00 | **0.002** | 0.66 | 0.28 – 1.04 | **0.001** |
| Observations | 98 | | | 98 | | |
| $R^2$ / $R^2$ adjusted | 0.097 / 0.088 | | | 0.110 / 0.100 | | |

### Supplementary datasets

**Dataset S1** Ploidy Survey.xlsx
**Dataset S2** Sample Metrics.xlsx
**Dataset S3** Selective Sweep Candidates.xlsx
**Dataset S4** Gene Ontology Enrichment.xlsx
**Dataset S5** *A_arenosa* $F_{ST}$ Window genes.xlsx
**Dataset S6** *C_amara* $F_{ST}$ Window genes.xlsx
**Dataset S7** Gene Ortholog Convergence.xlsx
**Dataset S8** Gene Homolog Convergence.xlsx
